# Blockade of redox second messengers inhibits JAK/STAT and MEK/ERK signaling sensitizing FLT3-mutant acute myeloid leukemia to targeted therapies

**DOI:** 10.1101/2022.03.09.483687

**Authors:** Zacary P. Germon, Jonathan R. Sillar, Abdul Mannan, Ryan J. Duchatel, Dilana Staudt, Heather C. Murray, Izac J. Findlay, Evangeline R. Jackson, Holly P. McEwen, Alicia M. Douglas, Tabitha McLachlan, John E. Schjenken, David A. Skerrett-Bryne, Honggang Huang, Marcella N. Melo-Braga, Maximilian W. Plank, Frank Alvaro, Janis Chamberlain, Geoff De Iuliis, R. John Aitken, Brett Nixon, Andrew H. Wei, Anoop K. Enjeti, Richard B. Lock, Martin R. Larsen, Heather Lee, Charles E. de Bock, Nicole M. Verrills, Matthew D. Dun

## Abstract

FLT3-mutations are diagnosed in 25-30% of patients with acute myeloid leukemia (AML) and are associated with a poor prognosis. AML is associated with the overproduction of reactive oxygen species (ROS), which drives genomic instability through the oxidation of DNA bases, promoting clonal evolution, treatment resistance and poor outcomes. ROS are also important second messengers, triggering cysteine oxidation in redox sensitive signaling proteins, however, the specific pathways influenced by ROS in AML remain enigmatic. Here we have surveyed the posttranslational architecture of primary AML patient samples and assessed oncogenic second messenger signaling. Signaling proteins responsible for growth and proliferation were differentially oxidized and phosphorylated between patient subtypes either harboring recuring mutation in FLT3 compared to patients expressing the wildtype-FLT3 receptor, particularly those mapping to the Src family kinases (SFKs). Patients harboring FLT3-mutations also showed increased oxidative posttranslational modifications in the GTPase Rac activated-NADPH oxidase-2 (NOX2) complex to drive autocratic ROS production. Pharmacological and molecular inhibition of NOX2 was cytotoxic specifically to FLT3-mutant AMLs, and reduced phosphorylation of the critical hematopoietic transcription factor STAT5 and MAPK/ERK to synergistically increase sensitivity to FLT3-inhibitors. NOX2 inhibition also reduced phosphorylation and cysteine oxidation of FLT3 in patient derived xenograft mouse models *in vivo*, highlighting an important link between oxidative stress and oncogenic signaling. Together, these data raise the promising possibility of targeting NOX2 in combination with FLT3-inhibitors to improve treatment of FLT3-mutant AML.

**One Sentence Summary:** FLT3-precision therapies have entered the clinic for AML however, their durability is limited. Here we identify the Rac-NOX2 complex as the major driver of redox second messenger signaling in FLT3-mutant AML. Molecular and pharmacological inhibition of NOX2 decreased FLT3, STAT5 and MEK/ERK signaling to delay leukemia progression, and synergistically combined with FLT3 inhibitors.

## Introduction

Mutations in the class III receptor tyrosine kinase (RTK) gene, *FLT3*, occur in approximately 25-30% of acute myeloid leukemia (AML) cases and are considered driver mutations (*1, 2*). These mutations result in constitutive activation of the receptor in the absence of FLT3-ligand, and include internal tandem duplications (ITD) of the juxtamembrane domain (*3*), and less commonly, point mutations within the tyrosine kinase domain (TKD) (*4, 5*). FLT3 inhibitors have recently been approved for use in FLT3-mutant AML patients, however improvements in outcomes are modest with relapse rates high (*2, 6*). There is growing evidence that reactive oxygen species (ROS) promote leukemogenesis (*7*), indeed, it is well recognized that ROS play a critical role in regulating normal hematopoiesis (*8, 9*). Oncogenic driver mutations are strongly linked to increased ROS production in myeloid malignancies (*10*). In AML, mutations in FLT3 appear to be most strongly linked to increased ROS production, however, mutations in the *RAS* oncogenes have also been implicated (*11–20*).

ROS are a heterogeneous group of molecules and radicals, previously thought to be by-products of cellular metabolism, that damage tissue and promote disease through DNA damage. In addition, it is now well recognized that ROS play an important role in cellular signaling in both physiological and pathological cellular processes (*21*). ROS regulate protein function via oxidation of the thiol functional group of cysteine residues (*22*). There are a wide range of known oxidative posttranslational modifications (oxPTMs) (*23*), which ultimately lead to alterations in protein structure and function. The two main cellular sources of ROS are the mitochondrial electron transport chain (ETC) and the NADPH oxidase (NOX) family; transmembrane proteins that reduce oxygen to superoxide via the transport of electrons across membranes (*24*). The NOX family is classified into p22^phox^-dependent (NOX1, NOX2, NOX3, NOX4) or calcium-dependent (NOX5, DUOX1 and DUOX2) enzymes (*24*). In FLT3-ITD leukemic cell lines, the FLT3-inhibitor PKC412 (midostaurin), the pan-NOX and flavoprotein inhibitor diphenyleneiodonium (DPI), and molecular knockdown of NOX1-NOX4 and p22^phox^ have each been shown to reduce ROS levels (*18*). p22^phox^ knockdown also decreased DNA double strand breaks in AML cell line models, thus supporting a role for NOX-derived ROS in driving genomic instability (*16*). The specific functional role of NOX enzymes in primary AML cells and their potential for therapeutic targeting, however, is not known.

Herein, we show that NOX2 and associated regulatory subunits are the key drivers of ROS production in primary FLT3-ITD AML and reveal that important tumor suppressors, oncogenic kinases and antioxidants all display increased cysteine oxidation in FLT3-ITD AML. By targeting NOX2 we show decreased oxPTMs in key regulatory enzymes including FLT3, GTPase Ras and NOX2, and show selective cytotoxicity in FLT3-ITD AML *in vitro*, and in primary FLT3-ITD patient-derived xenograft (PDX) mice. NOX2 inhibition also synergistically combined with FLT3 targeted therapies to increase the survival of a FLT3-ITD AML PDX mice.

## Results

### NADPH oxidase-2 drives oxidative posttranslational modifications (oxidome), enhancing oncogenic signaling in FLT3-ITD mutant AML

To gain a greater understanding of oncogenic signaling in FLT3-ITD AML, we employed quantitative analysis of the proteome, phosphoproteome and reversible oxidative posttranslational modifications of cysteines (hereafter the ‘oxidome’ or ‘oxPTMs’) to assess second messenger signaling, in three primary AML patient samples harboring FLT3-ITD mutations, and three primary AML patient samples wildtype for FLT3 (wt-FLT3) (Supplementary Table S1). In addition, one human FLT3-ITD AML cell line (MV4-11) was included, as were normal CD34+ bone marrow (NBM) cells (Fig. 1A). In completing these analyzes, we provide the first report of the AML oxidome; 2,946 proteins were identified as harboring reversibly oxidized posttranslational modifications of cysteines (oxPTM), and 2,219 proteins were identified as being phosphorylated, across all eight samples (Fig. 2A, Supplementary Fig. S1, Supplementary Table S2). A total of 962 proteins contained both phosphorylation and cysteine modifications with 60 individual peptides identified that harbored both phosphorylation and cysteine modifications.

**Fig. 1.**
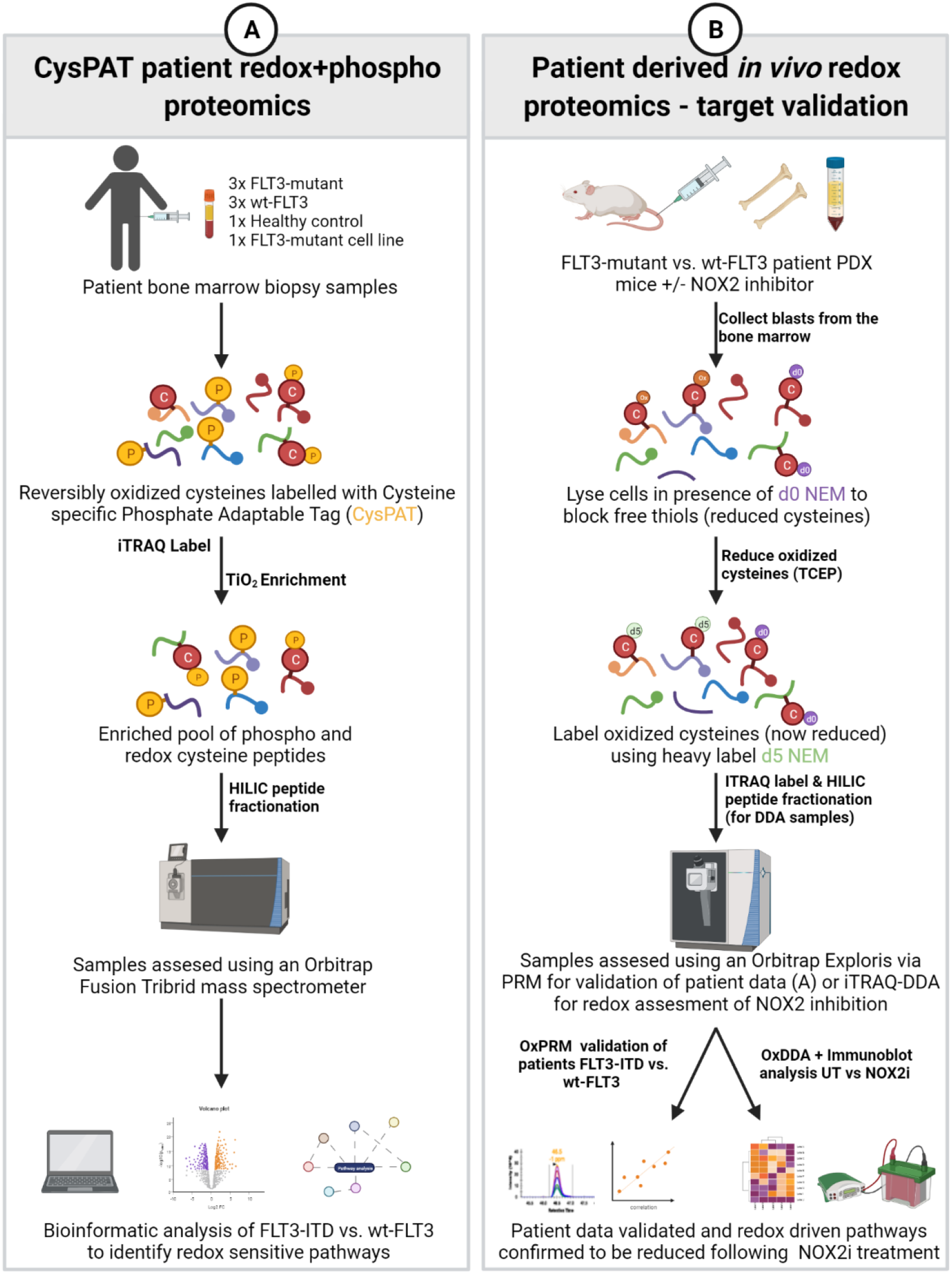
Redox proteomics workflows for the assessment of oxidative posttranslational modifications (OxPTMs) using AML patient samples *ex vivo* and patient derived xenograft models (PDX) *in vivo.* (**A**) Patient bone marrow trephine samples were collected at diagnosis and sequenced using a next generation sequencing (NGS) panel to identify driver mutations and 3x FLT3-ITD, 3x wt-FLT3, 1x normal bone marrow (NBM) control and 1x FLT3-ITD cell line were then subjected to CysPAT redox proteomics to assess phosphorylation and reversible oxidation of cysteines (oxPTMs). After free thiols were blocked with NEM, oxidized thiols were reduced and labelled with a Cysteine-specific Phosphonate Adaptable Tag (CysPAT) followed by iTRAQ labeling for relative quantification. Peptides containing reversibly oxidized cysteines (CysPAT) and phosphorylation were simultaneously enriched using TiO2 beads, subjected to HILIC fractionation and sequenced on an Orbitrap Mass Spectrometer. Bioinformatic analysis was performed to determine phospho- and oxPTMs changes in FLT3-ITD vs. wt-FLT3 patients. (**B**) Changes in oxPTMs were validated *in vivo* using primary patient samples and AML cell lines engrafted into NSG mice, either comparing FLT3-ITD vs. wt-FLT3 patient samples, or to assess how NOX2 inhibition modulates oxPTMs *in vivo*. Once leukemic burden reached 15-20% mice were treated with GSK2795039 or vehicle for 6 hrs then humanely culled (as per Institution ethics protocols). AML blasts isolated from the bone marrow were subjected to differential alkylation for the global assessment of oxPTMs (+/- NOX2 inhibition using iTRAQ) or using targeted proteomics using parallel reaction monitoring (PRM) to confirm primary patient oxPTMs identified in A. Cells were lysed in the presence of standard ‘d0’ NEM to alkylate free thiols. Then reversibly oxidized cysteines were reduced using TCEP and alkylated using a ‘heavy’ d5 NEM. Bioinformatics analyses and Western immunoblot was used to determine changes following NOX2 inhibition.

**Fig. 2.**
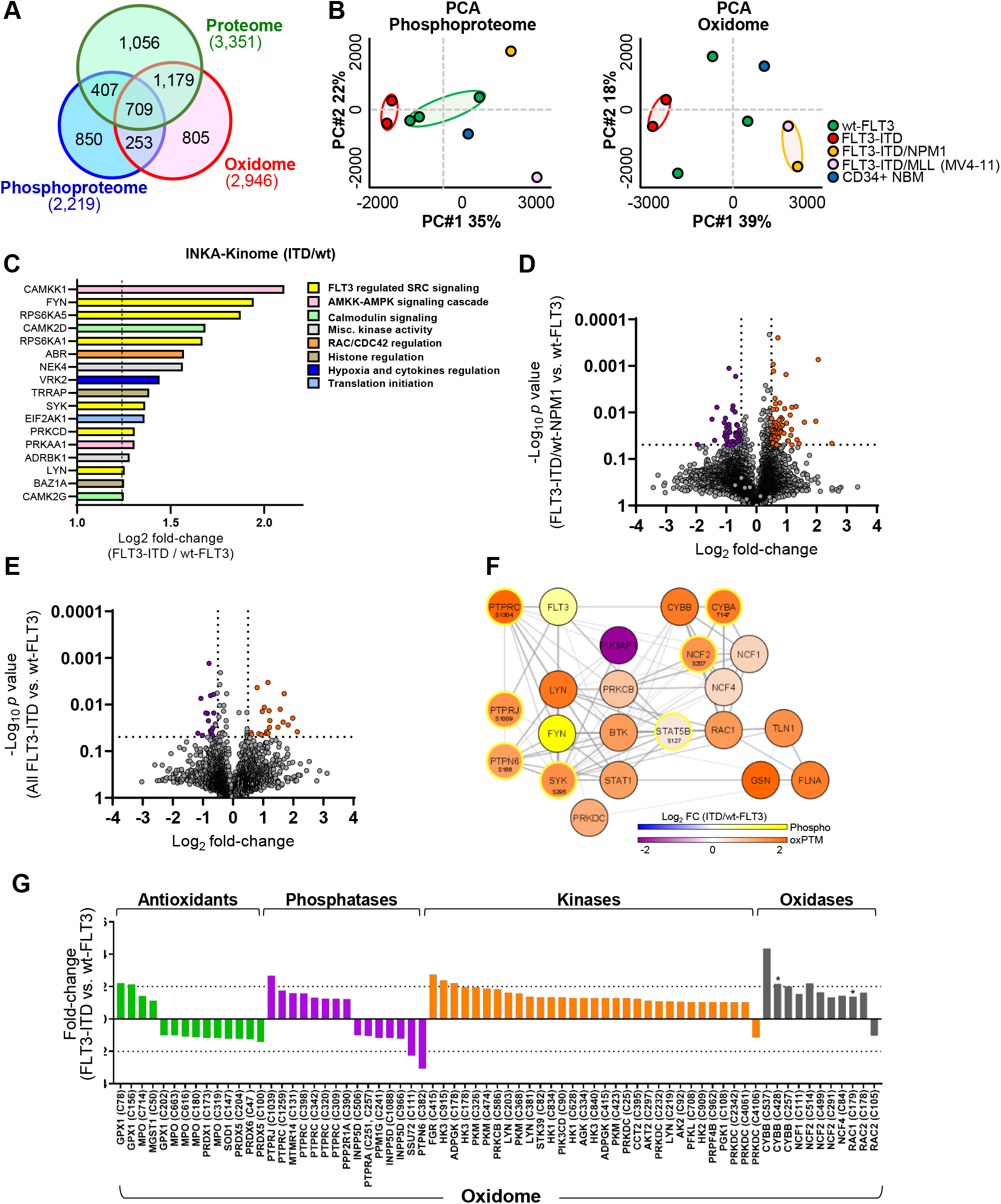
Global analysis of posttranslational modifications revealed redox activation of Rac-NOX2 and increased activity of the Src family kinases in FLT3-ITD+ primary AML patient samples. The proteome, phosphoproteome and cysteine-oxidome (oxidome) of six human AML blast samples, an AML cell line, and normal bone marrow control (NBM) were quantified using iTRAQ mass spectrometry. (**A**) Venn diagram analysis revealed 709 proteins harbored a non-modified, phosphorylated and cysteine oxidized peptide. 253 proteins harbored cysteine oxidized and phosphorylated peptides, while 407 proteins harbored a non-modified and phosphorylated peptide. 1,179 proteins harbored non-modified and cysteine oxidized peptides. (**B**) Principal component analysis (PCA) of the phosphoproteome and oxidome. (**C**) INKA (Integrative Inferred Kinase Activity) predicted activity of FLT3-ITD compared to wt-FLT3 phosphoproteomes. (**D**) Volcano plots comparing significantly regulated oxPTMs in patient blasts harboring FLT3-ITD/wt-NPM1 mutations vs. wt-FLT3; and (E) FLT3-ITD vs. wt-FLT3. (**F**) String database analysis of the top upregulated canonical pathways (FLT3-ITD compared to wt-FLT3) using proteins harboring oxPTMs (Orange/Purple fill = increased/decreased oxidation) including phosphorylation status (Yellow outline = increased phosphorylation) (**G**) Summary of the oxidome in all FLT3-ITD patients blasts compared to wt-FLT3 patient blasts. (* *p*<0.05 two-way ANOVA).

The phosphoproteome and oxidome were analyzed via principal component analysis (PCA) (Fig. 2B), with patients harboring FLT3-ITD mutations without Nucleophosmin 1 (*NPM1*) mutations (wt-NPM1) clustered in both analyses, whereas the total proteome analysis showed less consistent patient subtype clustering (Supplementary Fig. S1A,B). The phosphoproteomes were first interrogated using the ‘INKA’ pipeline (Integrative Inferred Kinase Activity) (*25*) to predict kinase (hyper) activity using multiple kinase- and substrate-centric modalities across the primary patient samples. Phosphoproteomes of FLT3-ITD patients’ blasts including those harboring NPM1 mutations were initially compared to the phosphoproteome of CD34+ NBM cells to eliminate kinase activity endogenous to hemopoietic stem and projector cell (HSC) signaling (Supplementary Table S3). Leukemia specific hyperactivated kinases were then compared between FLT3-ITD vs. wt-FLT3, with CAMKK1 and the Src-family kinase (SFK) FYN identified as the most hyperactivated kinases in FLT3-ITD samples (Fig. 2C). To assign molecular function to kinase activity specifically enriched in FLT3-ITD patient samples, hyperactivated kinases were analyzed by Reactome (*26*), revealing FLT3 regulated Src-signaling as the top ranked network in FLT3-ITD AML patient samples (Fig. 2C). Hyperactivated AMKK-AMPK signaling was the next top ranked kinase pathway in FLT3-ITD mutant AML; an intriguing result given that AMPK protects leukemia initiating cells (LICs) from metabolic and oxidative stressors (*27*). ABR kinase was also predicted to be hyperactivated, and when phosphorylated known to accelerate GTP hydrolysis of RAC1 or CDC42 to drive ROS production and drive oxidative stress (*28*), providing phosphoproteomic clues into the oxidative dysregulation of leukemia cells harboring kinase activating mutations (*9, 10*). INKA analysis of hyperactivated kinases in patient samples harboring wt-FLT3 showed a comparatively benign kinase signature, however also identified additional discrete components of AMPK signaling, and the core Hippo kinase STK4 (MST1) (*29*) as being hyperactivated (Supplementary Fig. S1D-F, Supplementary Table S3).

Unsupervised clustering of reversible cysteine oxidation showed conservation in oxPTMs in MV4-11 cells and the FLT3-ITD/NPM1 mutant primary sample, and a separate cluster of the two FLT3-ITD/wt-NPM1 patients (Supplementary Fig. 1G). To assess global differences between patient cohorts we first grouped FLT3-ITD/wt-NPM1 patients and compared oxPTMs with patients harboring wt-FLT3 using a log_2_ fold-change cutoff of ± 0.5. This analysis revealed 117 significantly regulated oxPTMs, in 111 unique proteins, of which 75 increased and 42 decreased in FLT3-ITD mutant patients compared to wt-FLT3 patients (Fig. 2D, Supplementary Table S4). Finally, stringent analysis of the oxidome using oxPTMs from all FLT3-ITD mutant patient samples compared to all wt-FLT3 patient samples, identified 25 significantly increased and 27 significantly decreased oxPTMs, corresponding to 46 unique proteins (Fig. 2E, Supplementary Table S4).

Significantly regulated oxPTMs in FLT3-ITD/wt-NPM1 patients compared to wt-FLT3 samples showed increased oxPTMs in the tyrosine kinases LYN-C203, SYK-C259 and BTK-C527, each known to be positively regulated by cysteine oxidation (Fig. 2E) (*10*), and like the phosphoproteome, act downstream of FLT3 (*30*). These data highlight the critical influence SFKs exert in the transmission of oncogenic messages in FLT3-ITD AML (*2*). Furthermore, RAC1/2 (C178/179) is known to be regulated by oxPTMs, and a potent activator of NOX2 (*31*). The subsequent increase in NOX2 activity further drives ROS production and potentially sets in train a cysteine oxidation feed-forward loop to exacerbate FLT3-ITD oncogenic signaling (Fig. 2F). While the functional significance is unknown, patients harboring wt-FLT3 AML showed increased oxPTMs in proteins regulating the spliceosomal cycle, including DExD-box helicase 39B (DDX39B-C165), and U2 small nuclear RNA auxiliary factor 2 (USAF2-C464), whereas U2 small nuclear RNA auxiliary factor 1 (USAF1-C67), and ATP-dependent RNA helicase (DDX42-C382) showed decreased oxPTMs (Supplementary Fig. S1≤ Supplementary Table S4).

To assess the global implications of oxPTMs in redox sensitive proteins in patients harboring FLT3-ITD mutations, we next compared oxPTMs with wt-FLT3 samples using Log_2_-fold 0.5 ± cutoff (Fig. 2G). Previous cell line studies showed PTPRJ (DEP-1) to be oxidized at the catalytic cysteine by elevated ROS, inactivating its phosphatase activity (*14*). Our study revealed a 2.6-fold increased oxPTM (C1039) in PTPRJ in FLT3-ITD AML samples compared to wt-FLT3 patients (Fig. 2G, Supplementary Table S5). When activated, PTPRJ has been shown to dephosphorylate FLT3 and thus negatively regulate its activity (*15*), although the direct effect on the activity of PTPRJ following oxPTM of C1039 is yet to be determined. Uniquely, our data also show that FLT3-ITD primary cells exhibit high levels of oxPTMs in the abundant hematopoietic transmembrane protein tyrosine phosphatase PTPRC, known as CD45 (Fig. 2F,G, Supplementary Table S5). PTPRC negatively regulates the activity of Src family tyrosine kinases (SFK) LYN, with PTPRC inactivation known to drive cellular transformation by selective and potent activation of the hemopoietic transcription factor STAT5 and the expression of genes necessary for proliferation, survival, and self-renewal (*32*). OxPTM of LYN at C466 in zebrafish promotes the activity of its downstream signaling pathways (*33*). This specific oxPTM was not identified in our studies, however, we found significantly increased LYN oxPTMs (C203 and C381) in the SH2 domain, and the kinase domain, respectively (Fig. 2G, Supplementary Table S4,S6). In addition, increased oxPTMs were observed in another SFK family member, FGR (C415), in FLT3-ITD vs. wt-FLT3 samples (Fig. 2F,G, Supplementary Table S6) highlighting the importance of oxidation of this tyrosine kinase family in oncogenic FLT3 signaling.

We next sought to explore whether antioxidant oxPTMs were differentially regulated between FLT3-ITD and wt-FLT3 AML. The oxidome analysis identified increased oxPTMs in GPX1 (C78 and C156) with a two-fold increased abundance in the FLT3-ITD samples, however the functional consequences of these modifications are not currently known (Fig. 2G, Supplementary Table S7).

### ROS production in FLT3-ITD AML is driven by increased oxPTMs in Rac-NOX2

Given the evidence supporting the ‘NOX family’ role in driving ROS production in AML, we sought to further characterize oxPTMs in the NOX isoforms and regulatory subunits in AML patient samples. Within the oxidome, all regulatory subunits of the NOX2 complex (RAC1, RAC2, CYBA/p22^phox^, NCF4/p40^phox^, NCF1/p47^phox^, NCF2/p67^phox^) were identified across all AML samples, without identification of any other NOX isoforms (Fig. 2G. Supplementary Table S8). Together, these proteome, phosphoproteome, and oxidome data (Fig. 2, Supplementary Fig. S1) suggest that NOX2 may be preferentially expressed and activated in FLT3-ITD AML. PTMs were identified in RAC1 (C179), RAC2 (C105, C178), p22^phox^ (T147), p40^phox^ (C84), p47^phox^ (C111), p67^phox^ (C291, C499, C514, S207) and NOX2 (C257, C428, C537) (Fig. 2F,G, Supplementary Table S8). All modified peptides showed increased abundance in FLT3-ITD AML samples apart from one oxPTM in RAC2 (C105), with RAC1 (C179) and NOX2 (C428) showing significantly increased oxPTMs in FLT3-ITD patients vs. wt-FLT3 patients. Phosphorylation of p22^phox^ (T147) showed a ∼2-fold increase in abundance in the FLT3-ITD samples and has previously been shown to enhance NADPH oxidase activity (*34*), whilst a novel phosphosite was identified in p67^phox^ (S207), which showed a ∼3.2-fold increase (Fig. 2G, Supplementary Table S8). Phosphorylation of p47^phox^ by protein kinase C (PKC) has been demonstrated upon activation of neutrophils by phorbol myristate acetate (PMA) (*35*). We observed increased oxPTM of p47^phox^ (C111) in FLT3-ITD samples compared to wt-FLT3 (Fig. 2G, Supplementary Table S8).

Searching against a database of known oxPTMs (RedoxDB) (*36*), only NOX2 (C537) has previously been reported to be functionally modified by cysteine oxidation and this has been proposed to increase NADPH oxidase activity (*37*). It is possible that the other modifications observed have functional impacts on NOX2 activity in FLT3-ITD AML that are yet to be characterized. We therefore sought to further characterize the role of NOX2 in driving leukemogenesis in FLT3-ITD AML by targeting the protein with small molecule inhibitors.

### NOX2 inhibitors reduce intracellular ROS and induce apoptosis through mitochondrial ROS production in FLT3-ITD AML

To determine the functional role and potential for therapeutic targeting of NOX2 in FLT3-ITD AML, we utilized three NOX inhibitors, VAS3947, GSK2795039 and APX115. The triazolo pyrimidine, VAS3947, has been shown to be a more specific NOX inhibitor compared to the historically used diphenylene iodonium (DPI), however, it does not have specificity for the NOX2 isoform (*38*). GSK2795039 is a small molecule inhibitor with unique specificity for NOX2 identified via high throughput screening and has been validated *in vitro* and *in vivo* (*39*). APX115 is an orally available pan-NOX inhibitor used in models of diabetic nephropathy and is currently in early phase II clinical trials (NCT04534439) (*40, 41*). All NOX inhibitors decreased cytoplasmic ROS production across AML cell lines highlighting the high-level of oxidative stress characterizing these cells (Supplementary Fig. 2A). However, NOX inhibition showed a commensurate increase in mitochondrial superoxide particularly in FLT3-ITD cells (Supplementary Fig. S2A). Apoptosis has previously been shown to increase the permeability of the mitochondrial membrane allowing ROS to be released into the cytoplasm (*42*). Further, mitochondrial ROS can trigger apoptosis; this apoptotic mechanism is a well characterized feature of arsenic trioxide in acute promyelocytic leukemia (*43*), and hypomethylating agents in AML (*44*). We next characterized the temporal changes observed in ROS and cell viability over a 24 hr period. An initial decrease in cytoplasmic superoxide was accompanied by a rise in mitochondrial superoxide in both the FLT3-ITD and wt-FLT3 cell lines, however, reduced cell viability was only seen in the cell lines harboring FLT3-ITD mutation (Supplementary Fig. S2B).

### NOX2 inhibitors and molecular interference of NOX2 results in selective killing of FLT3-ITD cell lines

We next sought to determine whether apoptosis was induced in AML cell lines using NOX inhibitors. FLT3-ITD cell lines were more sensitive to NOX2 inhibition with increased Annexin-V staining compared to wt-FLT3 cell lines (Fig. 3A,B). siRNA mediated knockdown of NOX2 resulted in a 48% and 43% decrease in NOX2 expression in MV4-11 (FLT3-ITD) and HL60 (wt-FLT3) cell lines respectively (Supplementary Fig. S2C), which led to increased apoptosis of FLT3-ITD cell lines compared to wt-FLT3 cells (Fig. 3A,C). These data confirm NOX2 dependence in FLT3 mutant AML (Fig. 3A-C) and reduce the possibility of ‘off-target’ effects mediating pharmacological NOX inhibition as a mechanism for the increased sensitivity of FLT3-ITD AML cell lines. Interestingly, wt-FLT3 cells also displayed sensitivity to APX115 (Fig. 3A,C, Supplementary Fig. S2D), which we postulate is due to its pan-NOX inhibitory effects (*40, 41*).

**Fig. 3.**
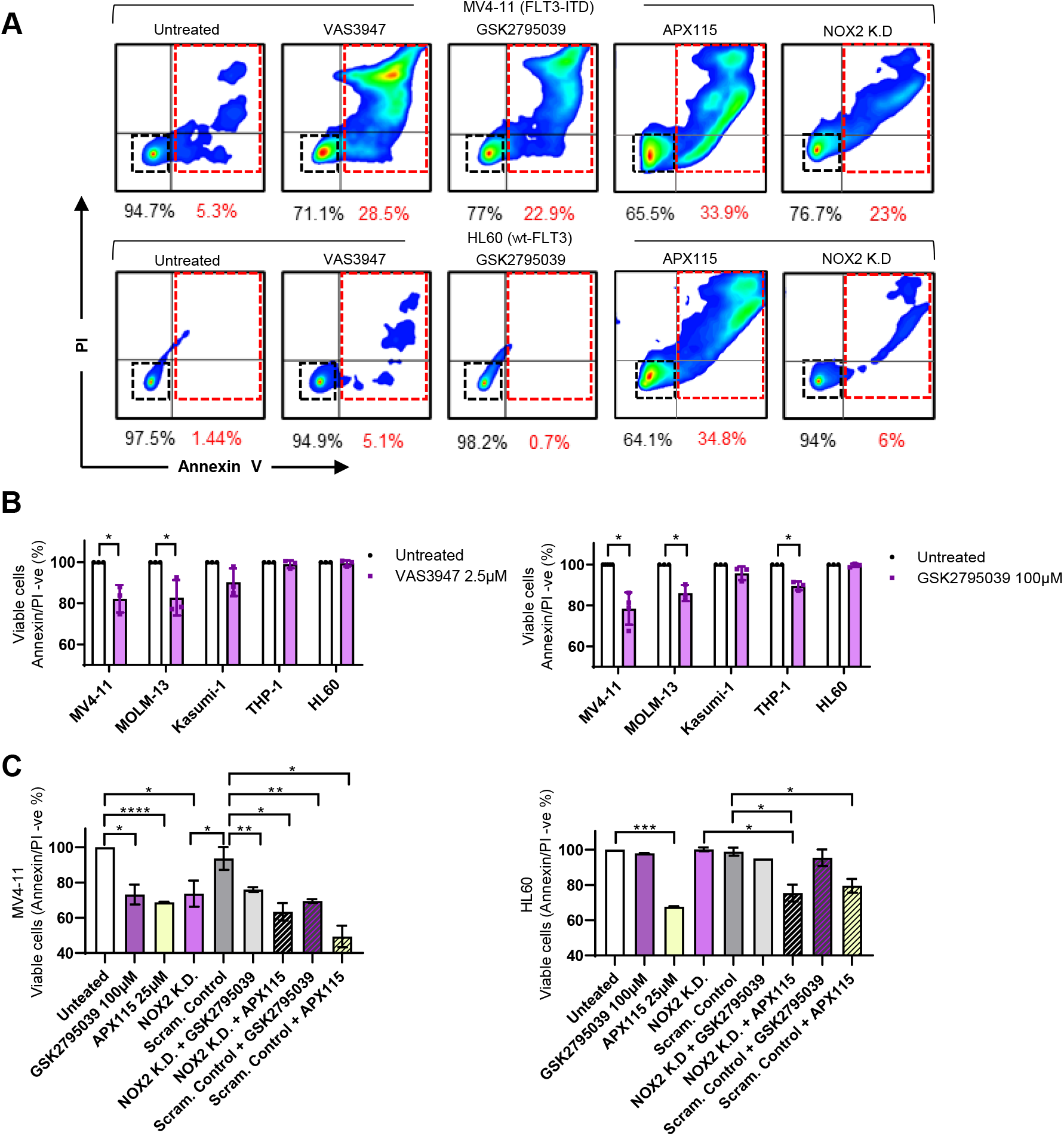
NOX2 inhibition induced apoptosis in human FLT3-ITD+ AML cell lines. (**A**) Representative images of FACS analysis +/- 24 hr treatment with 2µM VAS3947, 100µM GSK2795039, 25µM APX115 or siRNA knockdown of *CYBB* (NOX2) in MV4-11 (FLT3-ITD) or HL60 (wt-FLT3) human AML cell lines. (**B**) Annexin V staining of AML cells lines following 24-hr treatment with NOX inhibitors. (**C**) Quantitation of Annexin V positive MV4-11 (FLT3-ITD) and HL60 (wt-FLT3) cells following 24hrs treatment with NOX inhibitors +/- siRNA mediated knockdown of *CYBB* (NOX2) or scrambled control. Fluorescence was plotted relative to untreated control for each cell line, n=3 (\**p*<0.05, \*\**p*<0.01, \*\*\**p*<0.001, \*\*\**p*<0.0001, two-way Students T-Test, treatment vs. control).

### NOX2 inhibition in combination with FLT3-inhibition reduces oncogenic growth and survival signaling

We hypothesized that using NOX2 inhibitors to decrease ROS responsible for oxPTMs (Fig. 2) in combination with FLT3-inhibitors, would alter the activity of signaling pathways downstream of FLT3. To test this hypothesis, we first assessed expression of NOX2 and its regulatory subunits in AML cell lines using immunoblotting (Fig. 4A). NOX2, p22^phox^ and p47^phox^ showed increased expression in the FLT3-ITD cell lines (MV4-11, MOLM13) compared to wt-FLT3 cell lines (THP-1, HL60 and KAS-1). NOX2 protein expression was then correlated with sensitivity to the NOX2-specific inhibitor GSK2795039, or the pan-NOX inhibitor APX115 (Fig. 4A,B, and Supplementary Fig. S3A, respectively) which revealed a significant correlation between expression and sensitivity in AML cell lines where FLT3-ITD mutant cells lines with high NOX2 expression positively correlated with sensitivity (*p*=0.0429, Pearson correlation coefficient, *R^2^* = 0.678, Fig. 4B). No significant correlation was seen for FLT3-ITD mutant cells and sensitivity to APX115 (Supplementary Fig. S3A). Regulatory PTPs, SHP-1 and SHP-2 were expressed and tyrosine phosphorylated across our panel of cell lines (Supplementary Fig. 3B), however, STAT5, an important component of aberrant growth and/or anti-apoptotic signaling (*7, 8*), was uniquely phosphorylated at Y694 in FLT3-ITD mutant cells. JAK2 and LYN regulate STAT5 activity, with JAK2 phosphorylated at Y221 in FLT3-ITD (MV4-11) and c-KIT (KAS-1) mutant cell lines (Supplementary Fig. S3C). In contrast, the tyrosine kinase LYN which showed increased phosphorylation and oxPTMs at multiple sites in FLT3-ITD patient samples in the oxidome (Fig. 2C-G, Supplementary Table S2-S4,S6) was exclusively phosphorylated (Y507) in FLT3-ITD cell lines (Supplementary Fig. S3C).

**Fig. 4.**
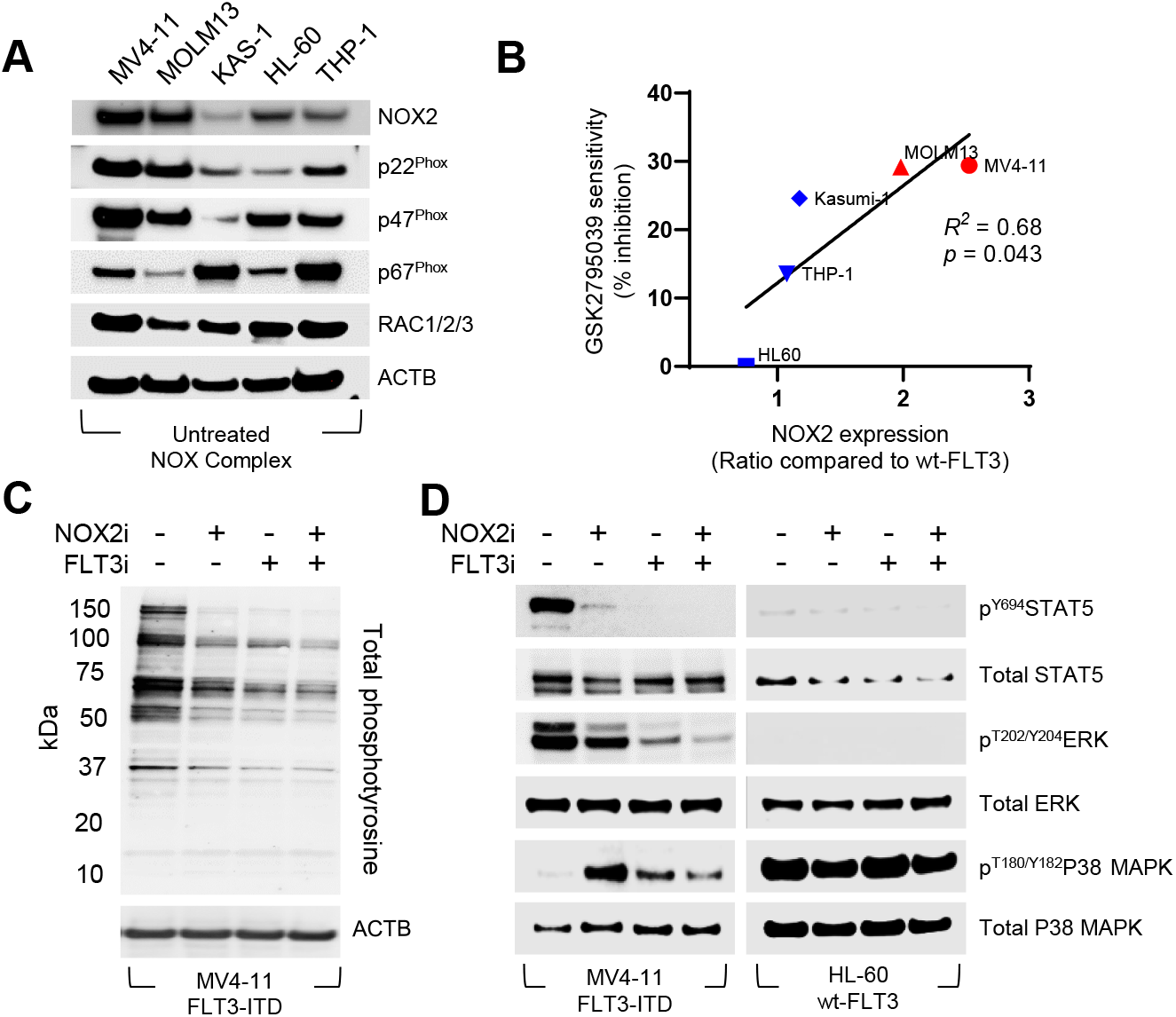
NOX2 expression is correlated with sensitivity to NOX2 inhibitors, reducing phosphotyrosine signaling in FLT3-ITD mutant AML. (A) Western immunoblotting expression analysis of proteins regulating the activity of NOX2 in a panel of human AML cell lines. (B) Pearson correlation analysis of NOX2 expression vs. sensitivity to NOX2 inhibition (GSK2795039) in AML cell lines (red = FLT3-ITD, blue = wt-FLT3). (C) Western immunoblot revealed reduced total phosphotyrosine signaling MV4-11 (FLT3-ITD) cell lines following NOX2 inhibition (GSK2795039) +/- FLT3 inhibition (quizartinib) for 90 min. (D) Western immunoblot revealed reduced phosphorylation STAT5, ERK and p38-MAPK in MV4-11 (FLT3-ITD), and not in HL60 (wt-FLT3) cells lines following NOX2 (GSK2795039) +/- FLT3 inhibition (quizartinib) (n=3 for each experiment, representative immunoblots are presented).

To assess whether we could modulate reversible oxPTMs, and hence the activity of redox sensitive enzymes, we treated FLT3-ITD cells with NOX2 inhibitors alone and in combination with the FLT3 specific inhibitor quizartinib (AC220). Following 90 min exposure, all treatments significantly reduced total phosphotyrosine levels, with the greatest reduction observed in the combination treatment group (Fig. 4C, Supplementary Fig. S3D). This prompted us to examine specific signaling proteins in FLT3-ITD (MV4-11) and wt-FLT3 (HL60) AML cell lines following NOX2 inhibition. Markedly, 90 min treatment with GSK2795039 ablated phosphorylation of STAT5 (Y694) alone and in combination with quizartinib (Fig. 4D). This approach also exclusively reduced phosphorylation of ERK (T202/Y204) in FLT3-ITD AML cell lines. Intrinsic defense mechanisms in healthy hematopoietic stem cells (HSCs) see the phosphorylation of p38-MAPK induce apoptosis following situations of increased oxidative damage, often a result of high ROS levels (*45*). p38-MAPK activity is silenced in many forms of cancer including FLT3-ITD AML, potentially promoting resistance to higher levels of ROS and, thus, gaining a survival advantage (*46*). After 90 min treatment with GSK2795039 alone, increased phosphorylation of p38-MAPK was observed, likely reflecting increased apoptosis (Fig. 4D). These results suggest that NOX2 is crucial for growth and survival of FLT3-ITD AML cells and highlights the link between NOX2 mediated redox signaling and the mutant receptor.

### NOX2 inhibition in combination with FLT3-inhibitors reduced proliferation of FLT3-ITD AML cell lines *in vitro* and *in vivo*

To determine the effect NOX2 inhibition plays on cellular proliferation in AML cell lines *in vitro* we first used FDC-P1 mouse myeloid progenitor cell lines harboring wt- and mutant-FLT3. A synergistic reduction of cell proliferation was observed when combining GSK2795039 and the FLT3-inhibitor quizartinib in cells harboring either wt-FLT3 stimulated with Flt ligand (FL) or constitutively active FLT3-ITD mutation but not in empty vector of control wt-FLT3 cells without FL (Fig. 5A, Supplementary Table S9). Human AML cell lines were treated with combinations of the NOX inhibitors VAS3947, APX115 or GSK2795039 and FLT3-inhibitors (sorafenib, midostaurin, quizartinib) which proved selectively lethal in FLT3-ITD cell lines (Fig. 5B, Supplementary Fig. S4A, Supplementary Table S9) and highly synergistic as defined by Chou-Talalay analysis (Fig. 5C). A moderate synergistic effect was also observed combining APX115 with sorafenib in a *RAS* mutant AML cell line (THP-1), although at higher concentrations than in the FLT3-ITD lines, and not with midostaurin or quizartinib which instead elicited an antagonistic effect (Fig. 5B,C, Supplementary Table S9). Similarly, there was no evidence of synergy in the wt-FLT3 or c-KIT mutant cell lines (HL60, Kasumi-1). The necessity of NOX2-driven ROS production for FLT3-ITD AML cell survival was further analyzed by assessment of Bliss Synergy (*47*), which showed GSK2795039 and APX115 when combined with midostaurin or sorafenib were highly synergistic in FLT3-ITD cell lines in contrast to HL60 cells (Fig. 5D). Similar results were observed by combining quizartinib with GSK2795039 (Supplementary Fig. S4A). Moderate synergy was observed via Chou-Talalay in the *RAS* mutant cell line THP-1 with the pan-NOX inhibitor APX115 and sorafenib, and an additive (Chou-Talalay) to moderate synergism (Bliss) was seen in the c-KIT mutant line Kasumi-1 with APX115 and midostaurin. As NOX2 was first identified in neutrophils, where it generates the respiratory burst required for pathogen inactivation (*24*), we assessed the effect of NOX2 inhibition alone and in combination with the multikinase inhibitor (including FLT3) sorafenib for 24hrs, on purified human neutrophils via an Annexin V assay, revealing no increase in apoptosis (Supplementary Fig. S4B).

**Fig. 5.**
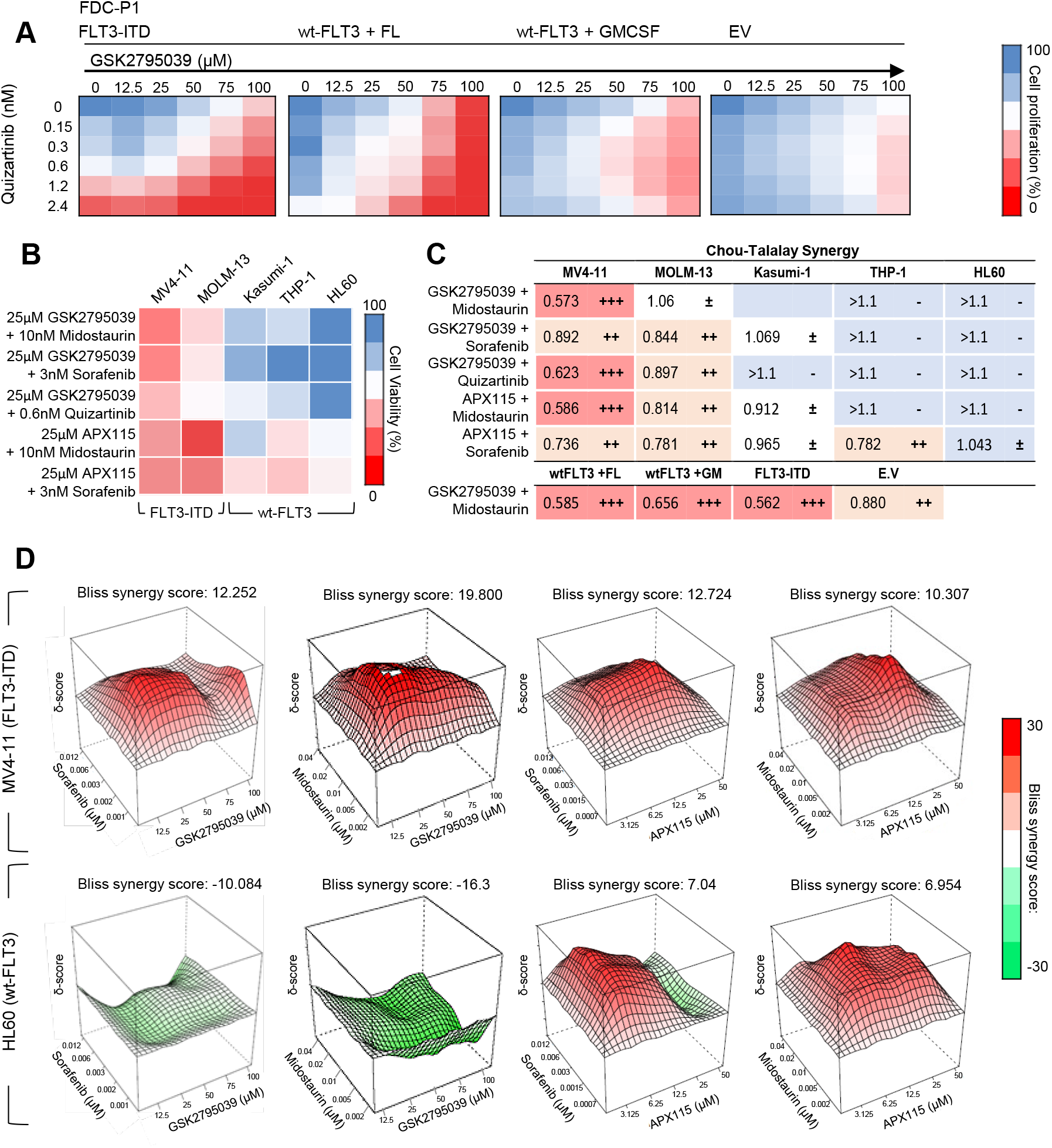
NOX2 inhibitors synergize with FLT3 inhibitors in cells signaling through FLT3. (**A**) Cell proliferation was assessed using the resazurin assay following 72-hr treatment with NOX2 inhibitor GSK2795039 and FLT3 inhibitor quizartinib in FDC.P1 cell lines transduced with wt-FLT3 (grown in the presence of FL or GMCSF), FLT3-ITD grown without ligand, or an empty vector (EV) grown in GM-CSF (cells generated as described (*76*)). (**B**) Cytotoxicity was determined using AML cell lines MV4-11 and MOLM13 (FLT3-ITD), HL60 and THP1 (mutant NRAS) and Kasumi-1 (mutant KIT) following NOX2 inhibition using GSK2795039 and APX-115 and FLT3 inhibition using quizartinib, sorafenib and midostaurin, (n=3 for each assay). (**C**) Combination index was calculated by Chou-Talalay analysis to determine synergy of NOX2 inhibitors GSK2795039 and APX115, with FLT3 inhibitors; - Antagonism (> 1.1), ± additive (0.9–1.1), ++ moderate synergism (0.7–0.9), +++ synergism (0.3–0.7). (**D**) Bliss synergy analysis predicting synergistic cytotoxicity (Bliss score >10) following treatment with midostaurin or sorafenib combined with GSK2795039 or APX115 in MV4-11 (FLT3-ITD), and an antagonistic (Bliss score <-10) or additive (Bliss score <10) effect in HL60 cells (wt-FLT3).

To assess the role the bone marrow microenvironment plays on the efficacy of NOX inhibition and hence the anti-AML potential of NOX2 inhibition in PDX mouse models of AML (Fig. 6A), NOD scid gamma (NSG) mice were engrafted with either a primary patient FLT3-ITD mutant or wt-FLT3 PDX (AML-16 or AML-5, respectively) (*48*). Following confirmation of engraftment via detection of human CD45+ cells (huCD45+) in the peripheral blood, mice were randomized and treated with GSK2795039. Response was tracked via flow cytometry of huCD45+ cells in peripheral blood (Supplementary Fig. S5A). After one week of treatment a significant reduction of huCD45+ % in the blood was identified in the FLT3-ITD cohort, whereas no significant difference was seen in the wt-FLT3 PDX (Fig. 6B, left panel). Survival was determined by extrapolating huCD45+ levels with a predetermined endpoint of 25% with the treated FLT3-ITD+ mice surviving significantly longer than the FLT3-ITD vehicle control group (12 days vs. 19 days, *p*=0.0002 Log-rank [Mantel-Cox] test) (Fig. 6B, middle panel). Notably, there was no difference in huCD45+ population, or survival, between vehicle and NOX2 inhibitor treated groups in the wt-FLT3 PDX (Fig. 6B right panel).

**Fig. 6.**
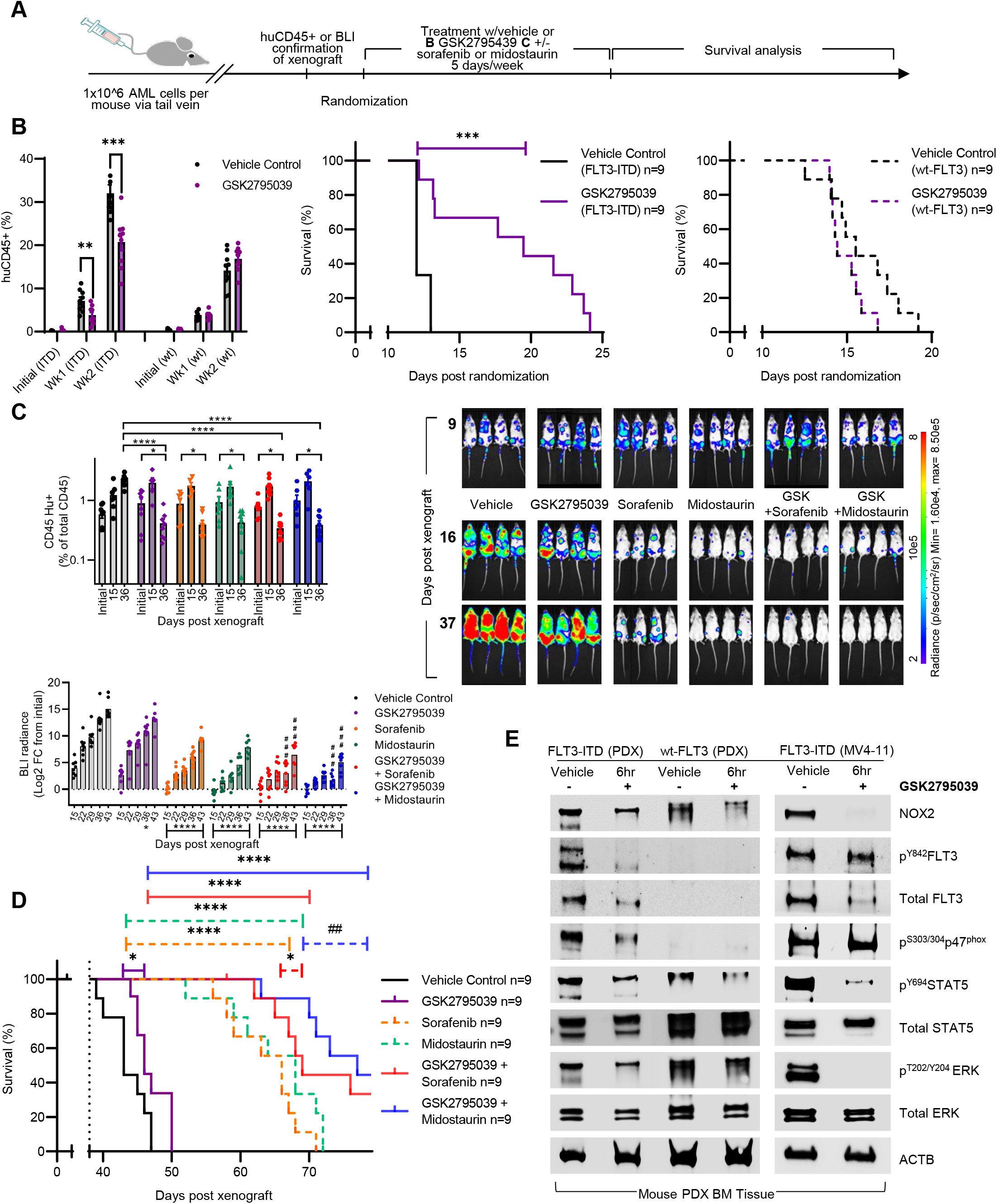
NOX2 specific inhibition using GSK2795039 increased survival of FLT3-ITD AML xenograft models and enhanced the durable response to FLT3 inhibitors. (**A**) Experimental timeline and key events. Leukemic burden was tracked weekly via huCD45+ flow cytometry of engrafted patient derived AML blasts (in **B**) or via *in vivo* bioluminescent imaging (BLI) for MV4-11-Luc+ cells (in **C,D**). Mice were randomized into treatment arms based on huCD45% in peripheral blood or BLI, with no significant difference between groups (Supplementary Fig. S5B). (**B**) Leukaemic burden was tracked by peripheral blood samples taken twice weekly and subjected to huCD45+ and msCD45+ FACs flow cytometry analysis and compared across groups +/- 100 mg/kg GSK2795039 treatment. Kaplan Meier survival analysis representing percentage survival of mice engrafted with AML patient derived cells (Log-rank (Mantel-Cox) test: *p*=0.0003 FLT3-ITD GSK2795039 vs. vehicle, or *p*=0.1147 wt-FLT3 GSK2795039 vs. vehicle). (**C**) MV4-11-Luc+ engrafted mice; peripheral blood samples were taken weekly and subjected to huCD45+ and msCD45+ by FACs analysis. Graph shows %huCD45+ plotted from initial reading (pre-treatment), after 1-week of treatment and at the end of 4-weeks treatment regime. Representative images of mice following 1-week of treatment (top) and at end of 4-weeks of treatment (bottom). Peak radiance tracked weekly plotted as a Log_2_ fold-change from the initial reading. (**D**) Kaplan Meier survival analysis representing percentage long-term survival following xenograft of MV4-11-Luc+ cells, end of treatment period represented by black dotted line. Kaplan Meier survival analysis revealed a significant survival advantage in mice treated with GSK2795039 (Log-rank (Mantel-Cox) test *p*=0.03 GSK2795039 vs. vehicle; *p*<0.0001 sorafenib vs. vehicle, *p*<0.0001 midostaurin vs. vehicle, *p=*0.01 GSK2795039 + sorafenib vs. sorafenib, *p=*0.002 GSK2795039 + midostaurin vs. midostaurin, synergism presented as ## determined as per Rose et al. (*77*). (**E**) Western immunoblot analysis of proteins isolated from bone marrow blasts of all three *in vivo* PDX models +/- 6 hr treatment with GSK2795039.

Given the success of NOX2 inhibition in our FLT3-ITD PDX model, we next tested the preclinical utility of combined NOX2 and FLT3-inhibition for the treatment of FLT3-ITD mutant AML in NSG mice engrafted with FLT3-ITD MV4-11-Luc+ cells (*49*). Once bioluminescence (BLI) reached a mean radiance of 3×10^6^ p/s, mice were randomized to receive vehicle, GSK2795039 (100 mg/kg), sorafenib (5 mg/kg), midostaurin (30 mg/kg) or GSK2795039 combined with either sorafenib (100 mg/kg GSK2795039 + 5 mg/kg sorafenib) or midostaurin (100 mg/kg GSK297039 + 30 mg/kg midostaurin) (Fig. 6A,C, Supplementary Fig. S5B). As a monotherapy, GSK2795039, reduced the proportion of leukemia cells in the peripheral blood, with BLI measurements also demonstrating a deeper reduction in leukemia burden after 4 weeks of GSK2795039 treatment (Fig. 6C), and significant survival benefit (Fig. 6D; 47 vs. 43 days for GSK2795039 vs. vehicle, *p*=0.03, Log-rank (Mantel-Cox) test). As expected, both sorafenib and midostaurin as monotherapies significantly increased survival; 66 (*p*<0.0001) and 68 days (*p*<0.0001) respectively, compared to the vehicle. The combination of GSK2795039 with sorafenib led to a significant survival benefit compared to the FLT3-inhibitor alone (73 days for GSK297039 + sorafenib, *p*=0.01), with the combination of GSK2795039 and midostaurin leading to both a significant and synergistic survival benefit compared to the FLT3-inhibitor alone (79 days for GSK2795039 + midostaurin, *p*=0.002) (Fig. 6D).

Bone marrow AML blast cells were harvested from both the PDX and MV4-11-Luc + models following acute inhibition of NOX2. Western blot of isolated blasts was performed and showed that GSK2975039 decreased FLT3 phosphorylation at Y842 located in the activation loop, revealing a direct link between NOX2-ROS and FLT3 activity (Fig. 6E). Downstream of FLT3, reduced STAT5 phosphorylation was also identified, with phosphorylation of ERK completely abolished using NOX2 inhibition alone, analogous to *in vitro* experiments. Notably, NOX2 abundance was decreased in both FLT3-ITD and wt-FLT3 PDX models, highlighting the NOX2 dependence of FLT3-ITD cells and the selectivity of GSK2795039. Phosphorylation of the NOX2 activating subunit p47^phox^ (S303/304) was also reduced, further indicating reduced NOX2 activation (Fig. 6E). These results demonstrate the interplay between NOX2 and ROS in leukemic blasts and the bone marrow microenvironment with the response in NOX2 active FLT3-ITD mutant samples remaining effective whilst, the limited benefit observed in the wt-FLT3 *in vitro* samples was now ablated *in vivo*. This is further highlighted by the significant increase in efficacy seen in the PDX bone marrow derived FLT3-ITD model compared to the cell line derived xenograft model, suggesting NOX2 inhibition to be effective in targeting leukemic blasts in the context of the bone marrow niche.

### Inhibition of NOX2 reduced FLT3 and RAC1/2 oxPTMs leading to decreased second messenger signaling *in vivo*

To determine the mechanisms underpinning the *in vivo* sensitivity of FLT3-ITD cells to NOX2 inhibition we performed global proteomic assessment of oxPTMs following acute treatment of FLT3-ITD xenograft models with GSK2795039 (Fig. 1B). Following GSK2795039 treatment, we identified 264 proteins containing oxPTMs, 61 of which were significantly modulated following GSK2795039 treatment (Fig. 7A, Supplementary Fig. 5C,D). Using these primary AML-PDX models we next validated the oxPTMs showing increased abundance in FLT3-ITD primary patient samples (Fig. 2). Targeted mass spectrometry was preformed using parallel reaction monitoring (PRM) using bone marrow blast cells isolated from primary FLT3-ITD and wt-FLT3 PDX models, showing FLT3-ITD patients harbor significantly increased abundance of oxPTMs in RAC1 (C179), RAC2 (C178) and a conserved peptide of RAC1 and RAC2 (C157) compared to wt-FLT3 (Fig. 7B).

**Fig. 7.**
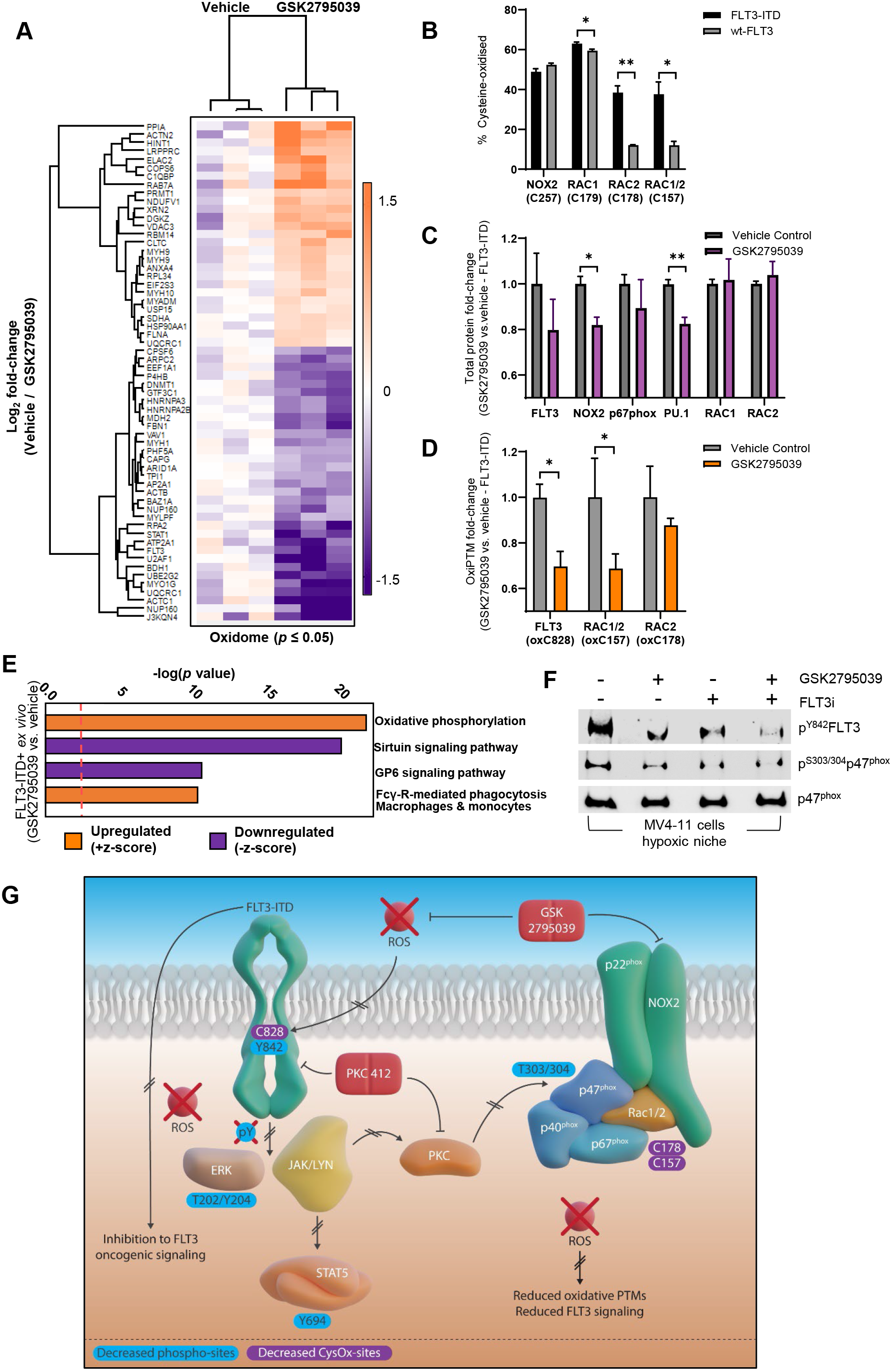
NOX2 inhibition modules redox homeostasis in FLT3-ITD AML blasts *in vivo* to enhance therapeutic benefit of FLT3-inhbitors. AML blast cells were isolated from PDX models following +/- 6-hr treatment with GSK2795039 and subjected to oxPTM analysis using iTRAQ proteomics. (**A**) Heatmap of significantly regulated (Log_2_-fold change 0.5 ±) reversibly oxidized cysteine proteins +/- 6 hr treatment with GSK2795039. (**B**) Validation of patient oxPTMs using parallel reaction monitoring (PRM) mass spectrometry using untreated blasts isolated from FLT3-ITD PDX engrafted mice vs. wt-FLT3 PDX mice. (**C**) Total proteome analysis in FLT3-ITD PDX mice following +/- 6 hr NOX2 inhibition with GSK2795039 represented as total protein fold-change compared to vehicle control. (**D**) Differential alkylation mass spectrometry analysis of oxPTM changes in key rac-FLT3-NOX2 peptides +/-6 hr GSK2795039 treatment. (**E**) Major pathways predicted to be modulated in FLT3-ITD vs wt-FLT3 PDX bone marrow blasts following *in vivo* treatment with GSK2795039 assessed by Ingenuity Pathway Analysis using significantly increased or decreased oxPTMs. (**F**) Western immunoblotting of the phosphorylation status of FLT3 and p47^phox^ in MV4-11 (FLT3-ITD) cells grown in hypoxic niche (<5% O_2_) +/- GSK2795039 alone or in combination with FLT3 inhibitor midostaurin. (**G**) Model of key oxPTM and phosphorylation events driving *in vivo* synergistic cytotoxicity induced by simultaneous inhibition of NOX2 and FLT3. The reduced superoxide production seen following NOX2 inhibition (GSK2795039) reduces oxPTMs and phosphorylation of FLT3-ITD in the activation loop, an event indispensable for FLT3-ITD mediated oncogenic signaling. NOX2 inhibition also reduces oxPTMs in the critical NOX2 activating protein Rac, thereby reducing the FLT3-NOX2 feedforward loop. Reduced activity of redox sensitive kinases through reduced superoxide production following NOX2 inhibition or inhibition of PKC using midostaurin, reduced phosphorylation of p47^phox^ in the NOX2 activation complex, a phosphorylation event that proceeds NOX2 activation. Combined inhibition of NOX2 and FLT3/PKC leads to complete loss of phosphorylation of p^Y842^FLT3 driving synergistic AML cell death *in vitro* and *in vivo*. Blue = phosphorylation site decreases following treatment. Purple = Cysteine oxidation site decreases following treatment.

Armed with this data, we assessed whether NOX2 inhibition in FLT3-ITD+ PDX samples would modulate oxPTMs. At the protein level, a significant *in vivo* decrease in NOX2 abundance corroborated results in Fig. 6E and supported the specificity of GSK2795039. *In vivo* NOX2 inhibition also significantly decreased protein abundance of PU.1 (Fig. 7C). Furthermore, proteins that showed increased oxPTMs in FLT3-ITD primary and *in vivo* patient samples (Fig. 2, Fig. 7B) were significantly reduced following NOX2 inhibition, including FLT3 (C828), RAC1/2 (C157) and RAC2 (C178) (Fig. 7D). IPA provided global insight into oxidome-related signaling changes following NOX2 inhibition *in vivo*. A shift in metabolic activity was seen with z-scores indicating a predicted increase in oxidative phosphorylation and decreased GP6 signaling, and reduced activity of Src family kinases FYN and LYN also predicted (Fig. 7E). MV4-11 cells grown in hypoxic conditions to mimic the bone marrow hypoxic niche, showed that midostaurin treatment alone and in combination with GSK2795039 reduced phosphorylation of FLT3 in the activation loop (Y842), and also decreased phosphorylation of PKC regulated p47^phox^ (S304) (Fig. 7F), an event that proceeds NOX2 activation (*50, 51*). ROS induced cysteine oxidation is indispensable for FLT3 driven oncogenic signaling (*19*), which we show is reversible with NOX2 inhibition (Fig. 7A-F), whilst activation of PKC requires cysteine oxidation for redox mediated complex formation with either SRC or LYN (*52*). This in turn leads to the phosphorylation of PKC and the activation of downstream signaling (*53*). Hence, inhibition of NOX2 using GSK2975039 leads to decreased phosphorylation of FLT3 and p47^phox^, analogous to treatment with midostaurin (Fig. 7F). The combination of midostaurin and GSK2975039 ablated phosphorylation of FLT3 and p47^phox^, helping to explain the *in vivo* therapeutic benefit of combined FLT3 and NOX2 inhibition for the treatment of FLT3-mutant AML (Fig. 7F,G).

## Discussion

Despite a rapid growth in knowledge of the genomic landscapes of AML, the condition remains a devastating disease with a poor prognosis (*54*). Among the alternative underlying etiologies that could contribute to this condition, ROS are emerging as key regulators of cellular signaling pathways, including those implicated in driving leukemogenesis (*9, 14, 19, 46, 55*). Thus, therapeutic manipulation of cellular ROS levels may hold promise as a novel treatment approach in AML (*10*). In keeping with this hypothesis, here we identified NOX2 as being preferentially activated in human primary AML blasts expressing the FLT3-ITD mutation. In addition, we confirmed increased reversible cysteine oxidation in proteins known to be regulated by ROS, including protein tyrosine phosphatases, Src family kinases and antioxidants in FLT3-ITD AML. It follows that pharmacological NOX2 inhibition led to a decrease in cytoplasmic superoxide, reduced activity of signaling proteins downstream of FLT3, as well as activation of apoptosis associated with mitochondrial ROS and p38 MAPK. Molecular knockdown of NOX2 led to increased apoptosis in FLT3-ITD cell lines as opposed to wt-FLT3 samples excluding off-target effects of the NOX2 inhibitors employed. Further, a synergistic anti-leukemic effect was observed by combining NOX2 inhibitors with clinically active FLT3-inhibitors in both AML cell lines and patient-derived xenograft mouse models.

In AML, mutations in *FLT3* and *RAS* are associated with increased ROS levels with most studies implicating the NOX family as the primary source (*11–13, 16*). More recently, in a cohort of 1069 patients, *FLT3*- and *RAS*-mutations correlated with high NOX2 (*CYBB*) expression and a 29 gene profile associated with metabolism that was able to predict poor survival, indicating NOX2 as a potential prognostic marker (*56*). In our studies herein, for the first-time using mass spectrometry in combination with cysteine specific enrichment, we have identified NOX2 and all regulatory subunits to be present in primary AML blasts. Furthermore, we show oxPTMs and phosphorylation in all proteins encompassing the NOX2-complex with increased abundance in FLT3-ITD AML, supporting increased activation in this kinase driven AML subset. In addition, we have shown increased reversible cysteine oxidation of key protein tyrosine phosphatases in FLT3-ITD AML, including PTPRJ, which has been previously reported (*14*). Although we recognize that the number of patients assessed was low, our sophisticated assessment of the posttranslational architecture of AML provides researchers with the tools to simultaneously assess second messenger and oncogenic signaling, clues which may help to reveal the mechanisms by which intracellular processes quickly adapt to therapeutic intervention using monotherapeutic approaches.

Importantly, NOX2 inhibition led to decreased phosphorylation of STAT5 and ERK, thus implicating NOX2 in downstream signaling pathways activated by FLT3. In fact, STAT5 has been shown to drive ROS production independent of JAK2 (*57*). Tyrosine phosphorylation of STAT5 drives interactions with RAC1, hence the inhibition of NOX2 and the subsequent reduction in FLT3 and RAC oxPTMs, potentially decreased STAT5 phosphorylation and therefore its interactions with RAC to create a negative feedback loop further reducing NOX2 activity. Indeed, inhibition of FLT3-ITD has been shown to decrease RAC1 activity and its binding to NOX (*10, 58, 59*). It has recently been demonstrated that RAC activation of NOX2 generates a feedforward loop. In a cell-free model both C157 and C18 oxidation within RAC was required for NOX2 activation (*31*). Our data demonstrate that NOX2 inhibition leads to decreased oxidation of C157, which would thus further reduce NOX2 activity.

Bohmer and colleagues have elegantly demonstrated that FLT3 signaling can be attenuated by replacement of critical cysteines. In addition to this, they demonstrated enhanced FLT3 activity with hydrogen peroxide stimulation (*19*). Using an *in vivo* model, and for the first time, we have demonstrated that NOX2 inhibition with GSK2795039 decreases oxidation of C828 adjacent to the activation loop of FLT3, which may, in turn reduce FLT3 activity. Of note, Bohmer and colleagues did not definitively prove that C828 was important for FLT3-ITD mediated cell transformation *in vitro*, however, it is likely that other cysteines would be reduced upon NOX2 inhibition, although they were not identified in our proteomic dataset. In addition, we have demonstrated decreased oxidation of C157 of RAC1/2 occurs following NOX2 inhibition.

NOX inhibition has previously been demonstrated to reduce intracellular ROS levels and cell proliferation in tyrosine kinase driven myeloid neoplasms, including chronic myeloid leukemia and FLT3-ITD AML cell lines (*60*). An increasing number of NOX inhibitors have been developed in recent years as the ‘NOX family’ emerges as a promising target in both malignant and non-malignant diseases (*10*). Although we recognize that GSK2795039 has limited clinical relevance due to its insolubility, it has been demonstrated to be specific for NOX2 both *in vitro* and *in vivo,* abrogating off-target effects through inhibition of other NOX isoforms by earlier compounds, such as VAS3947 or DPI (*39*) and hence, provided us with an ideal tool-compound to help reveal the dependence of FLT3-ITD AMLs on NOX2 activity. As previously discussed, it is possible that other kinase driven subsets of AML depend upon NOX2 derived ROS for a growth and survival advantage which in part may explain why our c-KIT mutant cell lines show some sensitivity to GSK2795039 *in vitro*. Further, inhibition of NOX-derived ROS may affect the tumor microenvironment. Tumor-associated macrophages produce ROS in a NOX-dependent manner, which results in NK and T cell dysfunction. Histamine dihydrochloride acting through H2 receptors is able to indirectly reduce NOX-derived ROS and protect NK and T cells from dysfunction and apoptosis in a paracrine manner (*61*). This has been tested in a Phase 3 clinical trial of post-consolidation therapy with histamine dihydrochloride and IL-2 in AML with improved leukemia-free survival observed in the experimental arm (*62, 63*). Further, NOX2-derived ROS was shown to stimulate bone marrow-derived stromal cells to transfer mitochondria to AML blasts via AML-derived tunneling nano-tubules, a process reversed using NOX2 knockout *in vivo* (*64*). These are examples in which ROS-high FLT3-ITD cells may gain a survival advantage by manipulating their environment. By disrupting these interactions with the microenvironment, we sensitized these cells. This in part explains why our FLT3-ITD PDX models derived directly from the bone marrow see an improved response *in vivo* to GSK2795039 compared to the cell-line derived xenograft which sees a modest response as a monotherapy. Furthermore, a role for NOX2 in regulating self-renewal of leukemic stem cells has recently been shown, with NOX2 knockout leading to impaired leukemogenesis in a murine model (*55*).

In summary, our data demonstrate that FLT3 mutant AML supports increased activity of NOX2, activation of tyrosine kinases, as well as inactivation of PTPs through reversible cysteine oxidation. NOX2 inhibition leads to reduced intracellular ROS, suppression of growth and survival pathways downstream of FLT3, and increased apoptosis associated with induction of mitochondrial ROS and restoration of p38-MAPK. Taken together, NOX2 has emerged as a novel target in FLT3-mutant AML with ongoing efforts to move drugs focused on this target into early phase clinical trials.

## Materials and Methods

### Materials

All chemicals used were purchased from Merck (Darmstadt, Germany), or Thermo Fisher Scientific (Rockford, IL) unless otherwise stated. Modified trypsin/Lys-C was from Promega (Madison, WI). Poros R2 and Poros Oligo R3 reversed-phase material were from Applied Biosystems (Forster city, CA). GELoader tips were from Eppendorf (Hamburg, Germany). The 3 m EmporeTM C8 disk was from 3 m Bioanalytical Technologies (St. Paul, MN). Titanium dioxide beads were purchased from GL Sciences Inc. (Tokyo, Japan). Alkaline phosphatase, PNGase F and endoprotease Asp-N were obtained from New England Biolabs (Ipswich, MA). Glyko® Sialidase C™ was from Prozyme (Hayward, CA). All solutions were made with ultrapure Milli-Q water (Millipore, Bedford, MA).

### Drugs

GSK2795039 was obtained from GlaxoSmithKline under a Materials Transfer Agreement (MTA) as was APX115 with AptaBio. Sorafenib was purchased from Selleckchem (Houston, TX); quizartinib and cytarabine from Cayman Chemical (Ann Arbor, MI); midostaurin from MedChemExpress (Monmouth Junction, NJ); VAS3947 from Merck Millipore (Burlington, MA) and hydrogen peroxide from Merck. Sorafenib, quizartinib, midostaurin, GSK2795039, VAS3947 and APX115 stock solutions were dissolved in DMSO. Cytarabine was resuspended in Milli-Q water at 50 mM stock concentration.

### Cell lines

The human AML cell lines MV4-11 and THP-1 were a kind gift from Dr. Kyu-Tae Kim (University of Newcastle, Callaghan, New South Wales, Australia); HL60 were a kind gift from Dr. Leonie K Ashman (University of Newcastle, Callaghan, New South Wales, Australia); and MOLM-13 were a kind gift from Dr. Jason Powell (Centre for Cancer Biology, Adelaide, South Australia, Australia). Cell lines were routinely screened for authenticity by the Australian Genome Research Facility. Normal CD34+ bone marrow mononuclear cells were purchased from Stemcell Technologies (Vancouver, BC, Canada) or Lonza (Basel, Switzerland). THP-1 and MOLM-13 cells were maintained in RPMI 1640 with 10% fetal calf serum (FCS), 2 mM L-glutamine and 25 mM HEPES with the addition of 0.05 mM ß-Mercaptoethanol for THP-1. MV4-11 and HL60 cells were maintained in DMEM with 10% FCS, 2 mM L-glutamine and 25 mM HEPES.

### Antibodies and Western blot analysis

Immunoblot analysis was performed using the following antibodies from Cell Signaling Technologies (Danvers, MA) (unless otherwise stated); NOX2 (Abcam, Cambridge, United Kingdom), p22^phox^ (Santa Cruz Biotechnology, Dallas, TX) p47^phox^, phospho^S303/304^p47^phox^ (Thermo Fisher Scientific, DE), p67^phox^ (Abcam), RAC1/2/3, phospho^y564^-SHP-1, total-SHP-1, phospho^Y542^-SHP-2, total-SHP-2, phospho^Y507^-LYN, total-LYN, phospho^Y694^-STAT5, total-STAT5, phospho-JAK2, total-JAK2, phospho^T202/Y204^-ERK, total-ERK, phospho^T180/Y182^-p38MAPK, total-p38MAPK, phospho^Y842^-FLT3, total phospho-tyrosine and β-actin (Sigma-Aldrich). Secondary antibodies were conjugated with horseradish peroxidase (Sigma-Aldrich).

Cells were lysed for Western blot analysis in ice cold RIPA buffer containing 5 mM Na_3_VO_4_, protease inhibitors cocktail and PhosSTOP (Roche, Penzberg, Germany) and sonicated for 2 × 10 s on ice, then mixed for 30 min at 4°C (as previously described (*65*)). Proteins were separated on NuPAGE Bis-Tris 4% to 12% gels (Invitrogen, Carlsbad, CA) and transferred onto 0.2 μm nitrocellulose membranes (Bio-Rad, Hercules, CA) for antibody staining (*66*). Bands were visualized via chemiluminescence using a ChemiDoc imager system (Bio-Rad).

### Primary AML patient blast proteomics

Proteins were purified from AML patients’ mononuclear cells as described (*49, 65*). AML blast cells were incubated in 1 ml of ice-cold 0.1 M Na2CO3 containing complete protease inhibitor (Roche, Penzberg, Germany) and phosphatase inhibitor PhosSTOP (Roche, Penzberg, Germany), (*67*) sonicated for 2 × 20 sec and incubated for 1 hr at 4 °C. The homogenates were then centrifuged at 100 000 × g for 90 min at 4 °C to enrich membrane and soluble proteins ( *68*) . Fractionated protein pellets were dried before being suspended in 6 M urea, 2 M thiourea, 2% SDS and alkylation of free thiol group of cysteines using 40 mM NEM (with the addition of complete protease inhibitor and phosphatase inhibitor PhosSTOP) (*69*). Lysates were loaded into 10 kDa spin filters to remove SDS and unreacted NEM, and protein concentration was subsequently determined via Qubit (*67*). The precipitated protein pellet was dissolved in 100 μL urea-buffer (6 M urea, 2 M thiourea), and reduced with 10 mM TCEP for 1 h at room temperature. The reduced protein was subsequently digested using Lys-C for 3 h. The solution was then diluted 8 times with 50 mM TEAB buffer to 0.75 M urea and 0.25 M thiourea, and trypsin (1:30) was added for further digestion at 37°C overnight. A total of 200 µg from each sample was labeled with iTRAQ 8plex reagents. Labeling efficiency was determined via MALDI-TOF/TOF MS. Samples were mixed 1:1 and stoichiometry determined once again via MALDI-TOF/TOF. The cysteine specific phosphonate adaptable tag (CysPAT) was synthesized as described (*70*). A multistage process of phosphorylated and cysteine oxidized peptide enrichment was achieved as previously described ( *71*) . Simultaneous enrichment for CysPAT labeled cysteine peptides and phosphorylated peptides was achieved using TiO2 as described (*70*). HILIC separated peptides were sequenced using an Orbitrap Fusion Tribrid Mass Spectrometer (Thermo Fisher Scientific, DE) coupled to an EASY-LC nanoflow HPLC system (Proxeon, DK). Samples were loaded onto in-house 2 cm pre-column packed with 3 µm Reprosil-Pur C18-AQ (Dr. Maisch GmbH, Germany) using an Easy-nLC II system (Proxeon, DK). The peptides were eluted from the pre-column onto an in-house packed Reprosil-Pur C18-AQ (17 cm x 75 μm, 3 μm; Dr. Maisch GmbH, Germany) column directly into the Orbitrap Fusion Tribrid Mass Spectrometer. The mobile phases were 95% acetonitrile (B buffer) and water (A buffer) both containing 0.1% formic acid. Peptides were eluted directly onto the analytical column using a gradient of 0% to 34% buffer B (90% acetonitrile, 0.1% formic acid) over 60 min. The Orbitrap Fusion Tribrid MS System was operated in full MS/data-dependent MS/MS mode. The Orbitrap mass analyzer was used at a resolution of 60000 to acquire full MS with an m/z range of 400-1400, incorporating a target automatic gain control (AGC) value of 2e^5^, and maximum fill times of 50 ms. The most intense multiply charged precursors (2-4 charges) were selected for higher-energy collision dissociation (HCD) fragmentation with a normalized collisional energy (NCE) of 40. MS/MS fragments were measured at an Orbitrap resolution of 15000 using an AGC target of 3e^4^, and maximum fill times of 100 ms.

Database searching of all .raw files was performed using Proteome Discoverer 2.1 (Thermo Fisher Scientific, DE). Mascot 2.2.3 and SEQUEST HT were used to search against the Swiss_Prot, Uniprot_Human database, (24,910 sequences, downloaded 10^th^ of February 2019). Database searching parameters included up to 2 missed cleavages to allow for full tryptic digestion, a precursor mass tolerance set to 10 ppm and fragment mass tolerance of 0.02 Da. Dynamic modifications included oxidation (M), phospho (S/T), phospho (Y), CysPAT (C), NEM (C) and iTRAQ-8plex. Interrogation of the corresponding reversed database was also performed to evaluate the false discovery rate (FDR) of peptide identification using Percolator on the basis of q-values which were estimated from the target-decoy search approach. To filter out target peptide spectrum matches (target-PSMs) over the decoy-PSMs, a fixed false discovery rate (FDR) of 1% was set at the peptide level ( *69*) . Subsequent analysis was conducted using ‘INKA’ (Integrative Inferred Kinase Activity) bioinformatics pipeline, IPA (Ingenuity Pathway Analysis), Reactome and Cystoscape using the (StringDB app). The mass spectrometry proteomics data have been deposited to the ProteomeXchange Consortium via the PRIDE partner repository (https://www.ebi.ac.uk/pride/login) with the dataset identifier PXD021995 and 10.6019/PXD021995”. **Username:** **Password:** RdL01g7j

### *In vivo* AML PDX proteomics

Bone marrow cells from human leukemia engrafted mice (MV4-11) were harvested following treatment with GSK2795039 or vehicle control. Cells were incubated in 900 µL of TUNES buffer (200 mM Tris, 6 M Urea, 100 mM NEM) for 1 hr at room temperature at 1 000 rpm to label free thiols. TCA was added to 20% v/v and precipitated proteins were then pelleted at 14 000 × g to remove NEM. Oxidized thiols were then reduced using TUNES buffer with NEM substituted for 10 mM TCEP. Differential alkylation was completed by labeling samples with 100 mM heavy labeled ‘d5’ NEM (Sigma-Aldrich) and protein concentration determined via Qubit (adapted from (*72*)). Samples were then digested and labeled using iTRAQ 8plex reagents as per patient sample proteomics above. HILIC separated peptides were sequenced using an Orbitrap Exploris 480 Mass Spectrometer (Thermo Fisher Scientific, DE) coupled to an EASY-LC nanoflow HPLC system (Thermo Dionex, Ultimate 3000 RSLC nano, Thermo Fisher Scientific). The peptides were eluted from the pre-column onto an Easy-spray (25 cm x 75 μm; Thermo Fisher Scientific, DE) column into the Orbitrap Exploris 480 mass spectrometer. The mobile phases were 95% acetonitrile (B buffer) and water (A buffer) both containing 0.1% formic acid. Peptides were eluted directly onto the analytical column over a 120 min gradient. The Orbitrap Exploris 480 MS was operated in full MS/data-dependent MS/MS mode. The Orbitrap mass analyzer was used at a resolution of 60,000 to acquire full MS with an m/z range of 360-1500, incorporating a target automatic gain control (AGC) value of 2e5, and maximum fill times of 50 ms. The most intense multiply charged precursors (2-4 charges) were selected for HCD fragmentation with a normalized collisional energy (NCE) of 36. MS/MS fragments were measured at an Orbitrap resolution of 15,000 using an AGC target of 3e4, and maximum fill times of 100 ms. Database searching of all .raw files was performed using Proteome Discoverer 2.5 (Thermo Fisher Scientific, DE). SEQUEST HT were used to search against the Swiss_Prot, Uniprot_Human database (97 512 sequences, downloaded 29^th^ of January 2021). Dynamic modifications included oxidation (M), phospho (S/T), phospho (Y), d5 NEM (C), NEM (C) and iTRAQ-8plex. Subsequent analysis was conducted using IPA (Ingenuity Pathway Analysis) assessing peptides with significant up or down fold-changes.

### Parallel reaction monitoring (PRM)

Peptides were injected onto a trapping column for preconcentration (Acclaim Pepmap100 20cm x 75 μm, 3μm C18, Thermo Fisher Scientific), followed by nanoflow LC (Thermo Dionex, Ultimate 3000 RSLC nano, Thermo Fisher Scientific). Peptide separation was achieved using a 15cm x 75µm, PepMap 3µm RSLC EasySpray C18 column (Thermo Fisher Scientific) with the following mobile phases: 0.1% formic acid in MS-grade water (solvent A) and 80% ACN combined with 0.1% formic acid (solvent B). Peptides were resolved using a 75-minute gradient that increased linearly from 2% to 35% solvent B, then ramped to 95% B with a constant flow of 400 nL/min. The peptide eluent flowed into a nano-electrospray emitter at the sampling region of a Q-Exactive Plus Orbitrap mass spectrometer (Thermo Fisher Scientific). The electrospray process was initiated by applying 2.20 kV to the liquid junction of the emitter, and data were acquired under the control of Xcalibur (Thermo Fisher Scientific) in PRM mode multiplexed two times. The precursors selected by PRM underwent high-energy collisional dissociation fragmentation with a normalized collision energy of 27.0, then measured by Orbitrap at a resolution of 17,500. Automatic gain control targets were 2E5 ions for Orbitrap scans. The raw MS data were processed using Skyline, version 21.2 (MacCoss Lab Software (*73*)). Inclusion list (Supplementary Table ST10).

### Detection of reactive oxygen species

Dihydroethidium (DHE) and MitoSOX™ Red reagent (Life Technologies, Australia) were used to detect intracellular cytoplasmic superoxide and mitochondrial superoxide, respectively. Briefly, cells were incubated with NADPH inhibitors (GSK2795039 or VAS3947), washed once in PBS and stained with dihydroethidium (DHE) or MitoSox Red reagent (adapted from (*74*)). After staining for 30 min cells were analyzed by FACSCanto flow cytometer (BD Biosciences, San Jose, CA). Data were analyzed using FlowJo software version 10.

### Cell proliferation and apoptosis

Cell viability was determined using a resazurin assay and cell death measured using the Annexin-V FITC apoptosis detection kit (BD Biosciences, San Jose, CA). Methods have been previously described (*65, 75*).

### Human AML xenograft models

All *in vivo* experimental procedures were conducted with approval from the University of Newcastle Animal Care and Ethics Committee (A-2017-733) and performed as previously described (*48, 49*). Engraftment levels were quantified by flow cytometry and expressed as the percentage of human CD45+ (hCD45+) cells to total hCD45+ and mouse CD45+ (mCD45+) cells in the tissue sample. Mice were given water and standard chow ad libitum. Eight-week-old mice were inoculated with MV4-11-luciferase cells (1×10^6^ cells, suspended in 100μL PBS) or AML PDXs (AML5 or AML16) (*48*) by injection into the lateral tail vein. Systemic leukemic burden in the MV4-11 model was assessed by bioluminescence imaging (BLI) using a Xenogen IVIS100 imager, following intraperitoneal injection of luciferin substrate (3mg/mouse; P1043 Promega). Leukemia burden in the peripheral blood was monitored twice weekly by flow cytometric analysis of hCD45+ proportions. Following erythrocyte lysis using ammonium chloride, white blood cells were stained with anti-human and anti-mouse CD45 antibodies, followed by analysis on a FACS Canto II flow cytometer (BD Biosciences 563879 and Biolegend 103115, respectively).

### Neutrophil extraction

Purified human neutrophils were extracted from whole blood using the EasySep™ Human Neutrophil Isolation Kit (Stemcell Technologies, Vancouver, BC) and performed in accordance with the manufacturer’s protocol.

### Statistical analysis

Graphs were produced using Graphpad Prism 7-9 software (La Jolla, CA, USA). Two sample paired and unpaired t-tests or two-way ANOVA was used to determine significant differences between groups except where otherwise indicated.

## Supplementary Materials

**Supplementary Figure S1.** Analysis of proteome, phosphoproteome and oxidome of AML patients.

**Supplementary Figure S2.** Assessment of ROS production, apoptosis and growth and proliferation of AML cell lines following molecular and pharmacological inhibition of NOX2.

**Supplementary Figure S3.** Assessment of NOX2 related proteins and sensitivity to NOX2 inhibitors.

**Supplementary Figure S4.** NOX2 inhibitors and FLT3 inhibitors combine to induce synergistic cell death in FLT3-ITD AML cell lines and are not cytoxic to purified human neutrophils.

**Supplementary Figure S5.** Assessment of leukemia burden of patient derived xenograft mouse models.

**Table S1.** Patient Samples Proteomics.

**Table S2.** Oxidome & Phosphoproteome.

**Table S3.** INKA Scores.

**Table S4.** Stat Significant PTMs -ITD+.

**Table S5.** Phosphatases.

**Table S6.** Kinases.

**Table S7.** Antioxidants

**Table S8.** NOX2 & Subunits PTMs

**Table S9.** Cytotoxicity IC50 Values.

**Table S10.** PRM in vivo PDX raw data.

## Acknowledgements

Mr. Nathan Smith from The University of Newcastle Analytical and Biomolecular Research Facility (ABRF) provided MS support. The Academic and Research Computing Support (ARCS) team, within IT Services at the University of Newcastle, provided high performance computing (HPC) infrastructure for supporting the bioinformatics. Graphical abstract and Fig. 1 created using www.biorender.com. Summary cartoon in Fig. 7 created with thanks to www.somersault1824.com. GlaxoSmithKline provided the NOX2 inhibitor GSK2795039 and AptaBio provided pan-NOX inhibitor APX115.

## Author Contributions

Contribution: J.R.S., Z.P.G., M.R.L., N.M.V. and M.D.D., conceived and designed the study and interpreted the results; Z.P.G., J.R.S., A.M., R.D., H.C.M, I.F., M.R.L. and M.D.D., conducted the experiments and performed data analysis; D.S., I.J.F., H.C.M., J.E.S., D.S.B., M.N.B., H.H., and C.E.d.B. helped with experimental work and/or interpretation of results; D.S.B., assisted bioinformatic analyzes; M.W.P., G.D.I, B.N., R.J.A., F.A. and J.C., provided discipline specific expertise; A.K.E. and A.W., assisted with obtaining and processing of patient samples; J.R.S., Z.P.G., A.M.D and M.D.D., wrote and edited the manuscript; and all authors discussed the results and commented on the manuscript.

## Data and materials availability

MTA agreements and publically accessible data as described in the materials and methods.

## Graphical Abstract

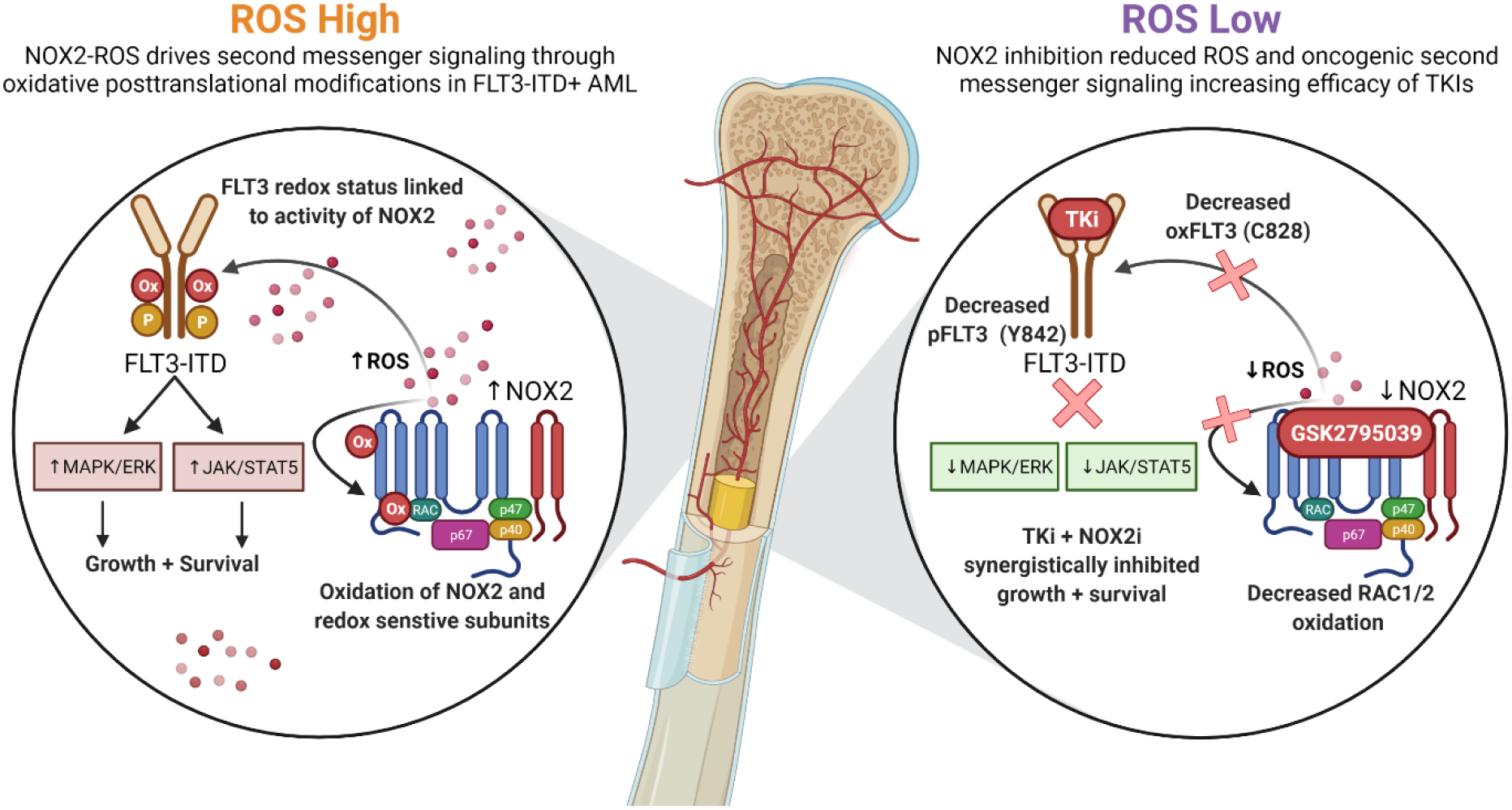

---

**Supplementary Fig. S1.**
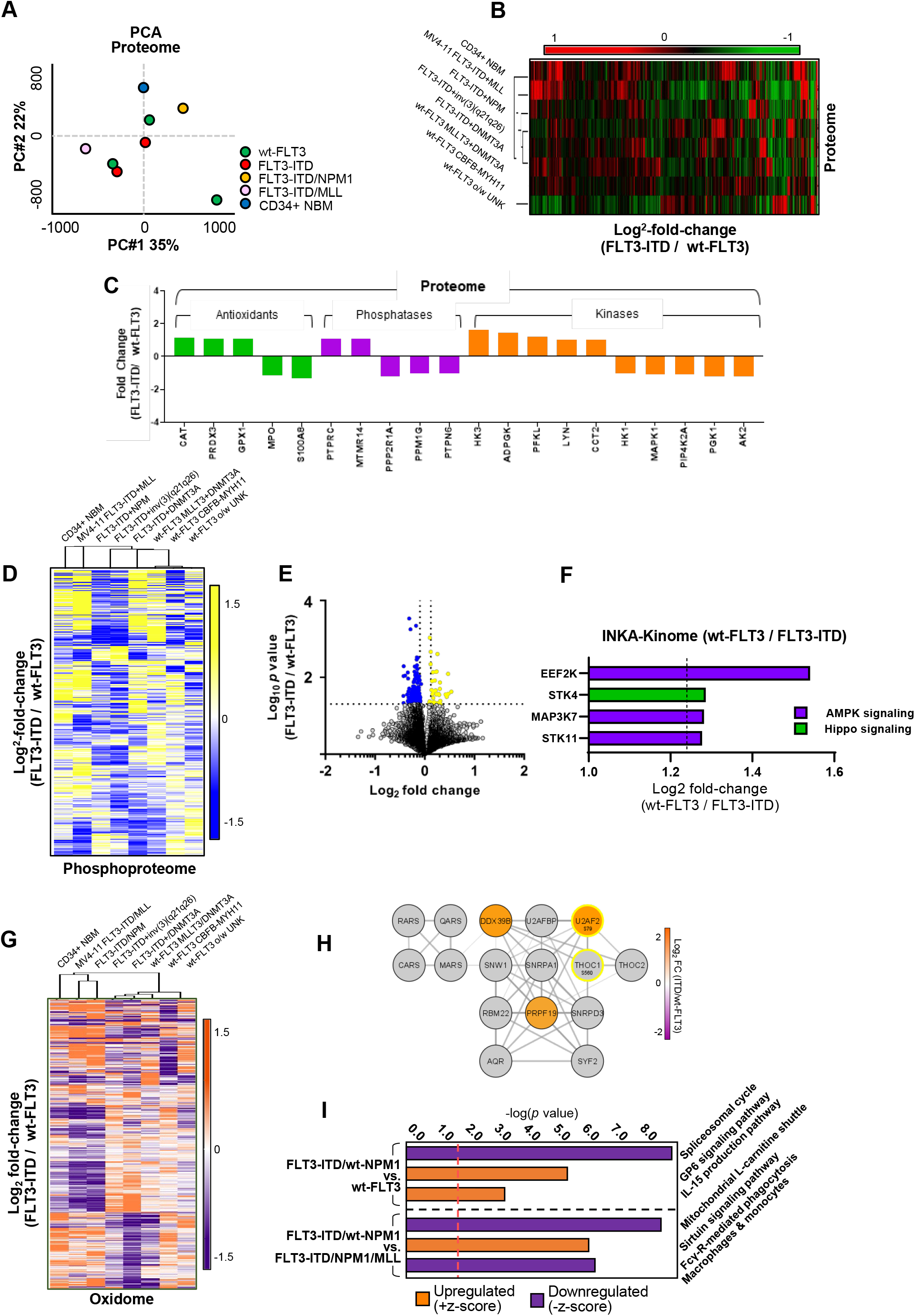
Analysis of proteome, phosphoproteome and oxidome of AML patients. (**A**) Proteome PCA clustering analysis. (**B**) Heatmap clustering of individual patient samples and corresponding somatic mutations. Heatmap displays the proteome as Log_2_ fold-change of each sample compared to the average of wt-FLT3 samples. (**C**) Summary of redox sensitive proteins in the data set revealed no significantly increased or decreased abundance of these key regulatory and redox sensitivity proteins. (**D**) Heatmap clustering of the phosphoproteome represented as a fold-change ratio (FLT-ITD/wt-FLT3). (**E**) Volcano plot of phosphopeptides plotting Log_2_ fold-change of FLT3-ITD compared to wt-FLT3 samples. Yellow/Blue represent significantly up/down regulated peptides. (**F**) INKA predicted activity of wt-FLT3 vs. to FLT3-ITD phosphoproteomes. (**G**) Heatmap clustering oxPTMs of individual patient samples and corresponding somatic mutations. (**H**) String database analysis of the top upregulated canonical pathways (wt-FLT3 vs. to FLT3-ITD) using proteins harbouring oxPTMs (Orange/Purple = increased/decreased oxidation) including phosphorylation status (Yellow = increased phosphorylation) (**I**) IPA analysis of significant proteins regulated by oxPTMs.

**Supplementary Fig. S2.**
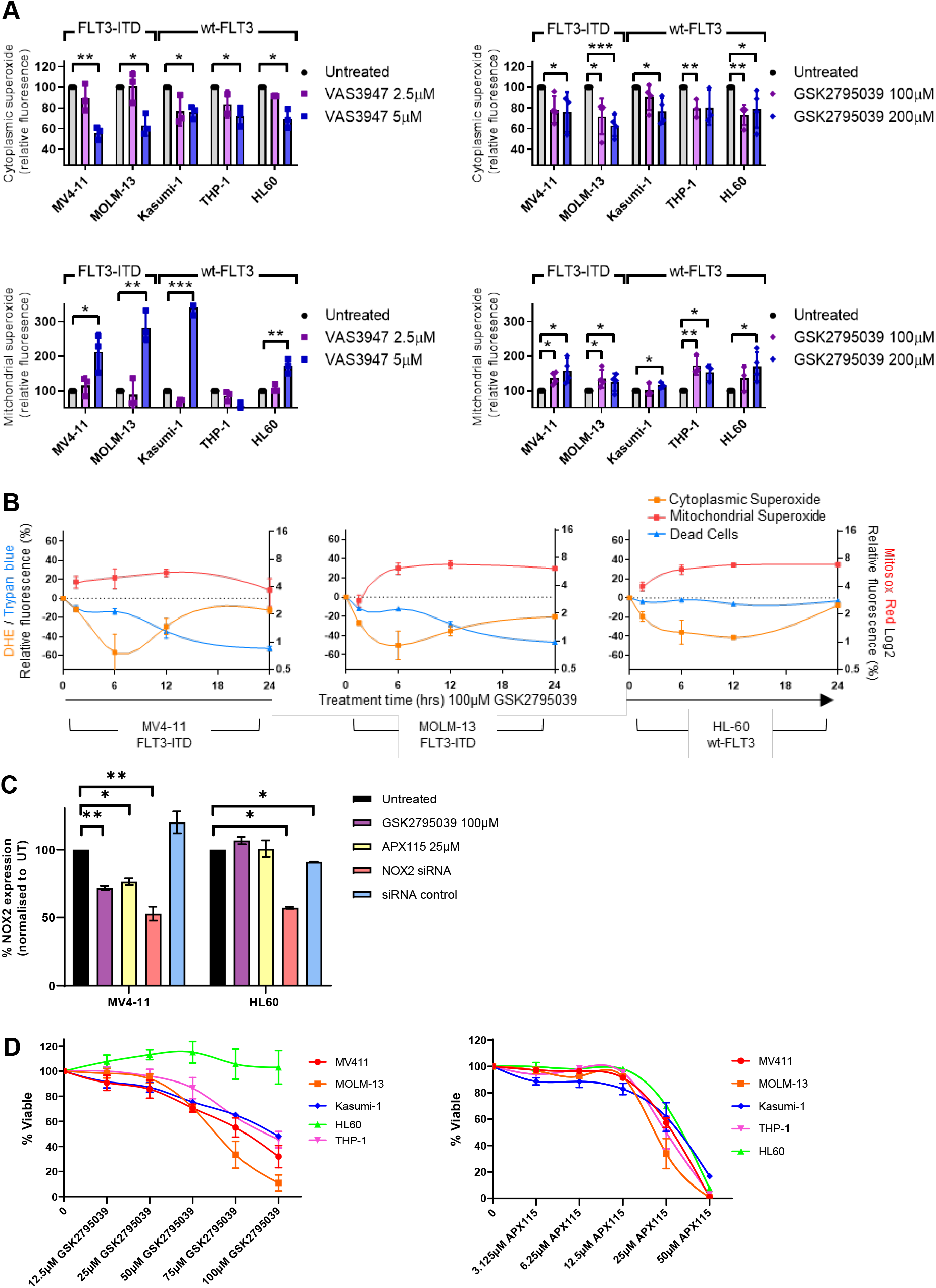
Assessment of ROS production, apoptosis and growth and proliferation of AML cell lines following molecular and pharmacological inhibition of NOX2. (**A**) Cytoplasmic (top) or mitochondrial (bottom) superoxide was quantified via flow cytometry using DHE or MitoSox red fluorescent probes applied to a panel of human AML cell lines +/- 1 hr treatment with NOX inhibitors. (**B**) MV4-11, MOLM-13 and HL-60 cell lines were treated with NOX2 inhibitor GSK2795039 for 90 min, 6 hr, 12 hr and 24 hr, cells stained with DHE (cytoplasmic superoxide – orange), Mitosox red (mitochondrial superoxide – red) and Trypan blue (dead cells – blue). (**C**) Flow cytometry measurement of NOX2 expression following: 72 hr NOX2 siRNA knockdown (*CYBB*/NOX2 and scrambled control), NOX inhibitor GSK2795039 or APX115. NOX2 expression is normalized to untreated samples. (**D**) Cell proliferation assay measured by Resazurin fluorescence in a panel of human AML cell lines following 72 hr treatment with increasing doses of NOX inhibitors GSK2795039 or APX115 (n=4).

**Supplementary Fig. S3.**
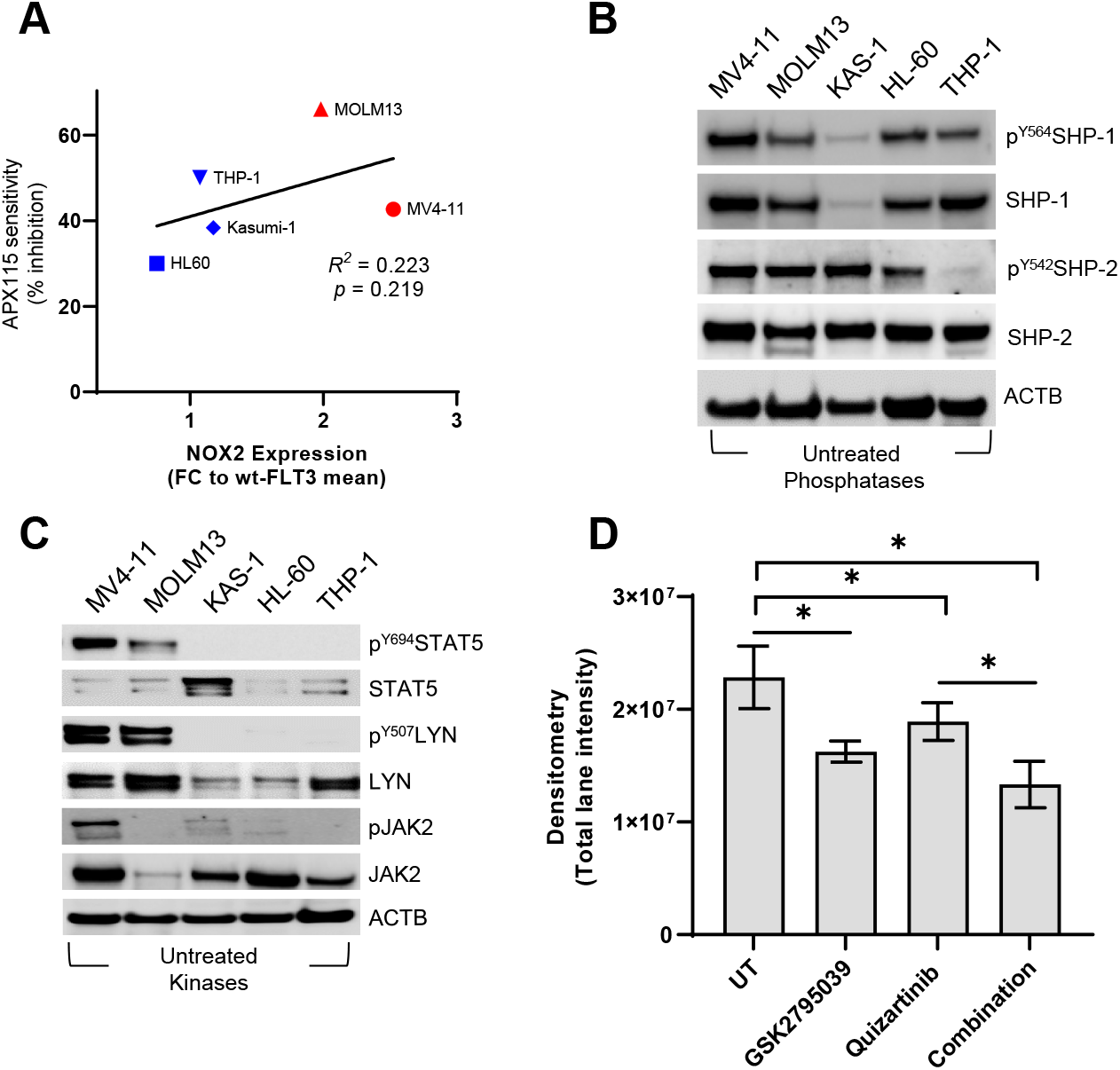
Assessment of NOX2 related proteins and sensitivity to NOX2 inhibitors. (**A**) NOX2 expression across AML cell lines stratified by FLT3-ITD mutational status (Red = FLT3-ITD, Blue = wt-FLT3) determined via densitometry and correlated with sensitivity to the pan NOX inhibitor APX115. Western immunoblot analysis (**B**) tyrosine phosphorylation of protein tyrosine phosphatases, (**C**) tyrosine phosphorylation of kinases, and **(D**) densitometry quantified changes in total phosphotyrosine signaling following NOX2 inhibition, FLT3 inhibition, or by the combination of both MV4-11 cells (n=3). Representative blot presented in main text Fig. 3C (\**p*<0.05, Two-way Students T-Test).

**Supplementary Fig. S4.**
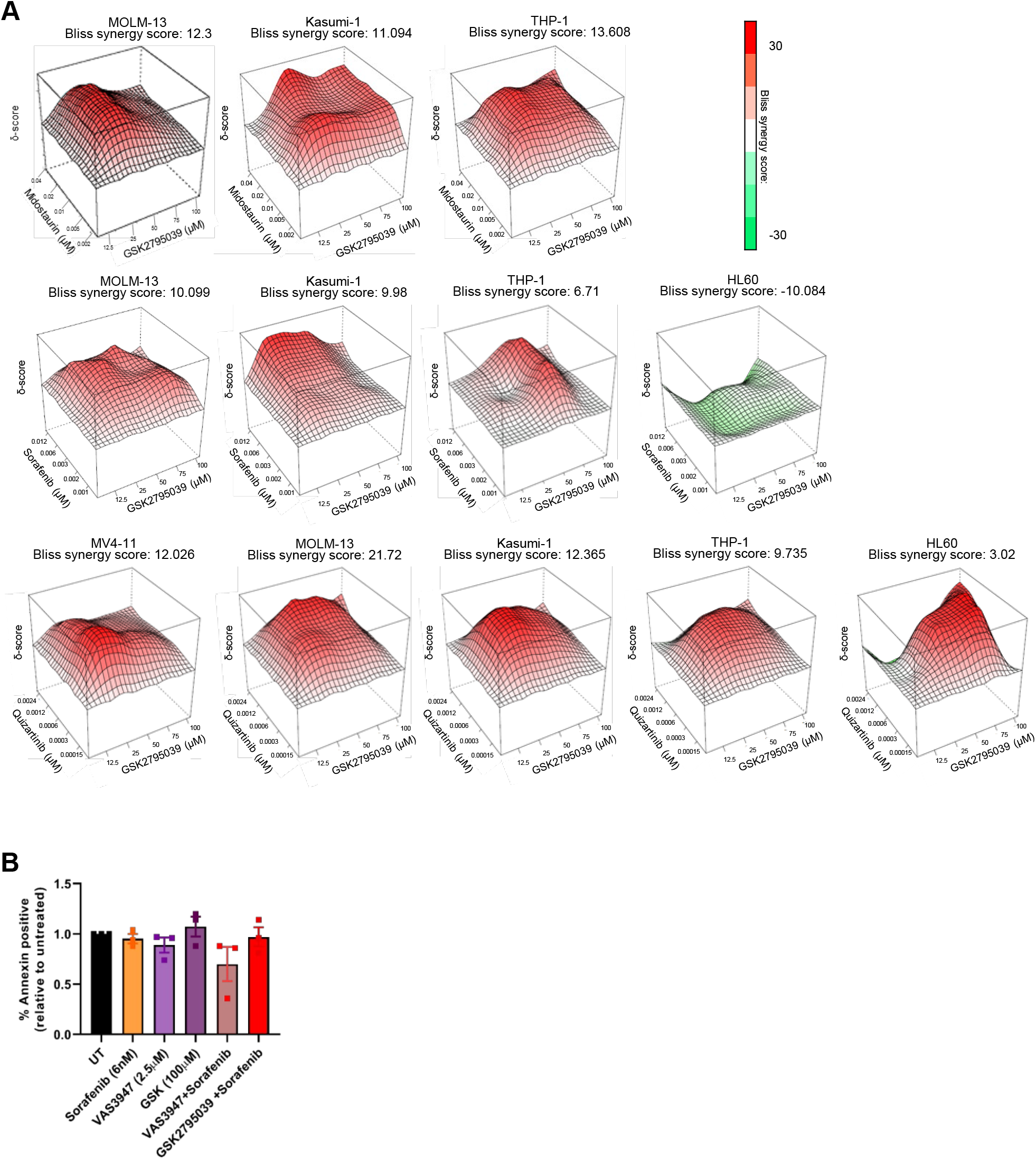
NOX2 inhibitors and FLT3 inhibitors combine to induce synergistic cell death in FLT3-ITD AML cell lines, without causing cell death of purified human neutrophils. (**A**) Bliss synergy analysis showing synergistic cytotoxicity (Bliss score >10) or antagonism (Bliss <-10) using NOX2 inhibitors in combination with midostaurin, sorafenib or quizartinib in MV4-11, MOLM13, Kasumi-1, THP-1 and HL60 human cell lines. Scores calculated based on n=3 replicates of resazurin proliferation assays. (**B**) Blood samples were taken from healthy patients and neutrophils purified via the EasySep™ Human Neutrophil Isolation Kit followed by treatment with VAS3947 (2.5µM) and GSK2795039 (100µM) both alone and in combination with Sorafenib (6nM). Samples were then stained with Annexin V and propidium iodide and analyzed via flow cytometry to determine apoptosis and cell death compared to untreated. No significant difference was observed in treated samples compared to untreated (n=3 patients were used).

**Supplementary Fig. S5.**
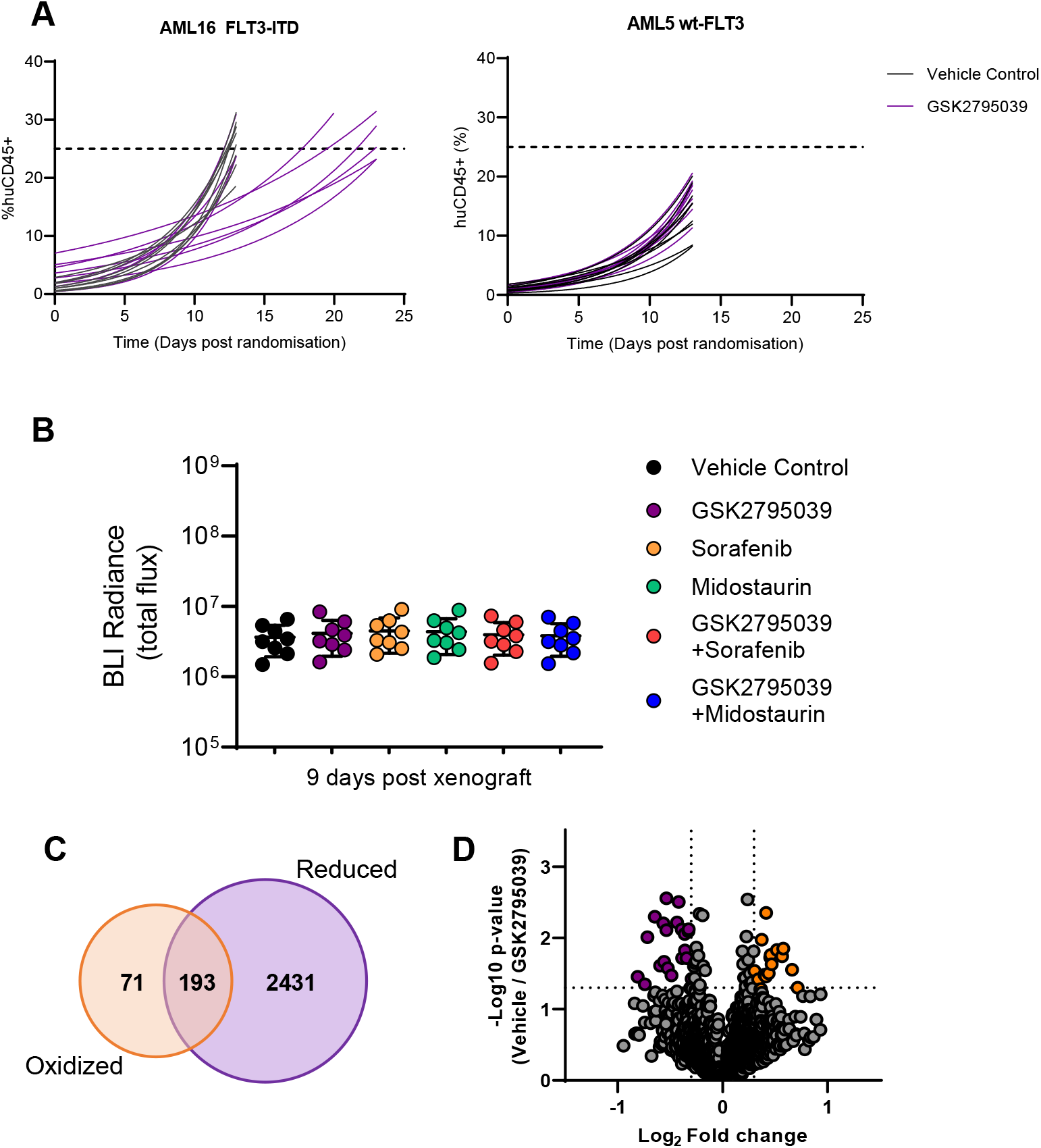
Assessment of leukemia burden of patient derived xenograft mouse models. (**A**) Percentage huCD45+ cells detected via flow cytometry in the peripheral blood of patient derived xenograft mice plotted bi-weekly. Data are plotted and fitted with an exponential growth curve to extrapolate the time point at which mice reach 25% huCD45 in the peripheral blood (pre-determined ethical endpoint). Left represents FLT3-ITD PDX engrafted mice treated with a vehicle control (black) or 100mg/kg GSK2795039 (purple) and right represents wt-FLT3 PDX engrafted mice. (**B**) Nine days after engraftment (1 × 10^6^ Luciferase expressing MV4-11 cells), bioluminescence was used to determine the level of engraftment in each mouse. Mice were then randomized into treatment arms with 9 successfully engrafted animals in each group and total radiance plotted. No significant difference observed between groups (one-way ANOVA). Bone marrow cells were harvested following 6 hr treatment with either 100mg/kg GSK2795039 (n=3) or vehicle (n=3), followed by sequential labeling of reduced and oxidized cysteines. Peptides were labeled with iTRAQ tags, subjected to HILIC fractionation, and sequenced via high resolution mass spectrometry (**C**) Proteins identified represented in a Venn-diagram where 2 624 proteins harbored reduced cysteines, 264 with reversibly oxidized cysteines and 193 proteins containing cysteines in both states. (**D**) Proteins were mapped on a volcano plot plotting a Log_2_ fold-change of GSK2795039 vs. vehicle treated samples (Log_10_ p-value).

