## Supplementary Figures for "Blockade of redox second messengers inhibits JAK/STAT and MEK/ERK signaling sensitizing FLT3-mutant acute myeloid leukemia to targeted therapies"

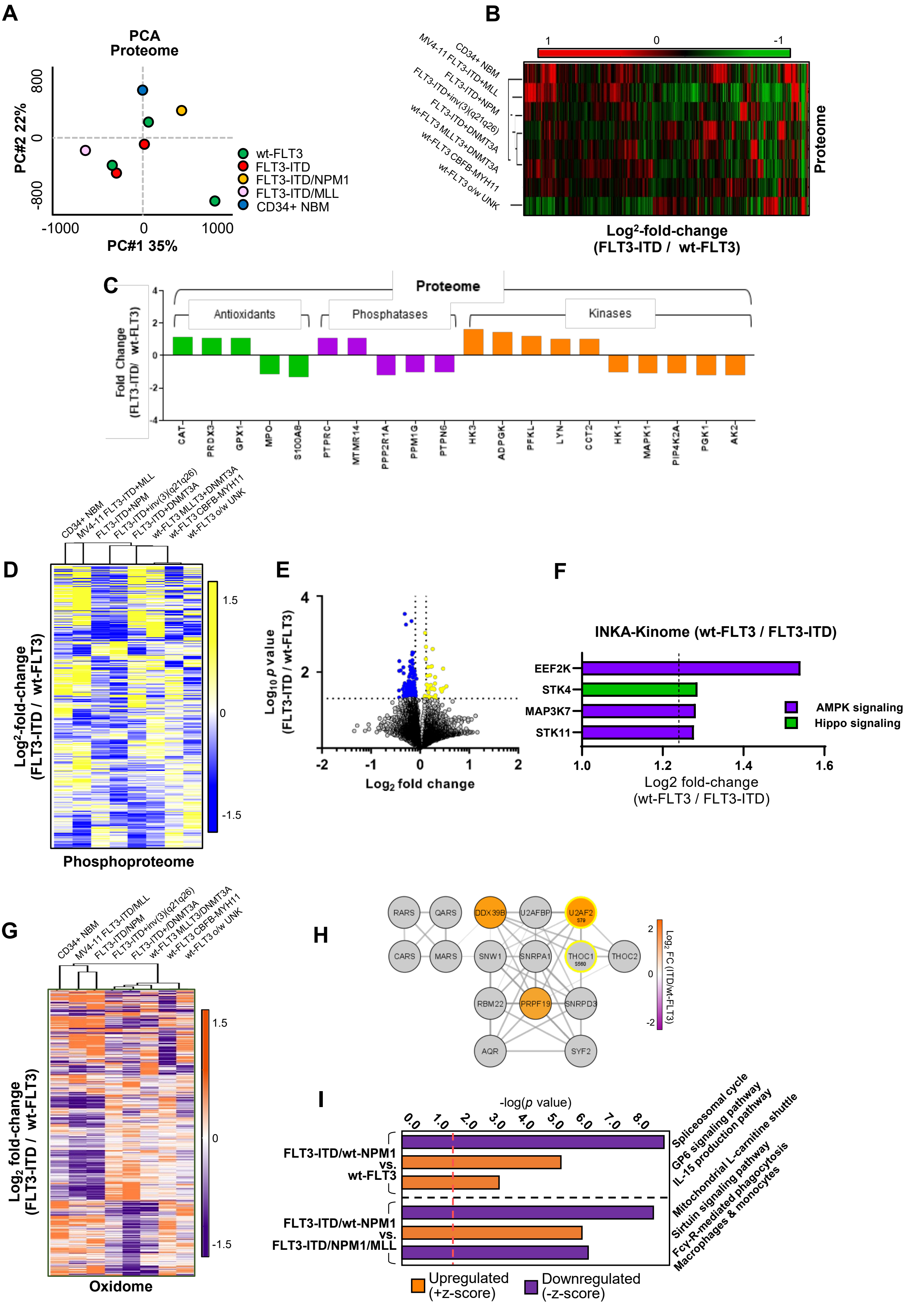


**Supplementary Figure S1.**

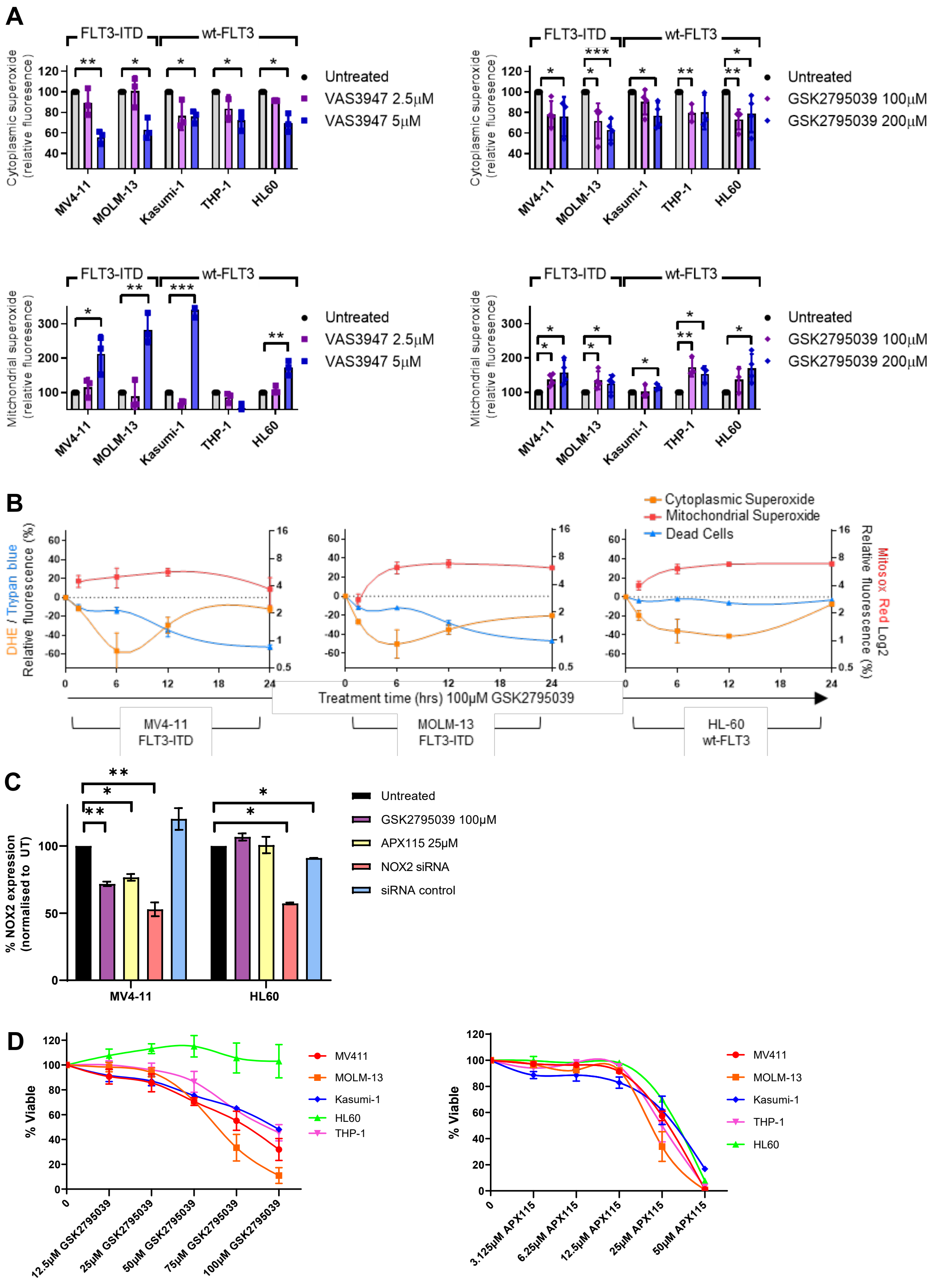


**Supplementary Figure S2**

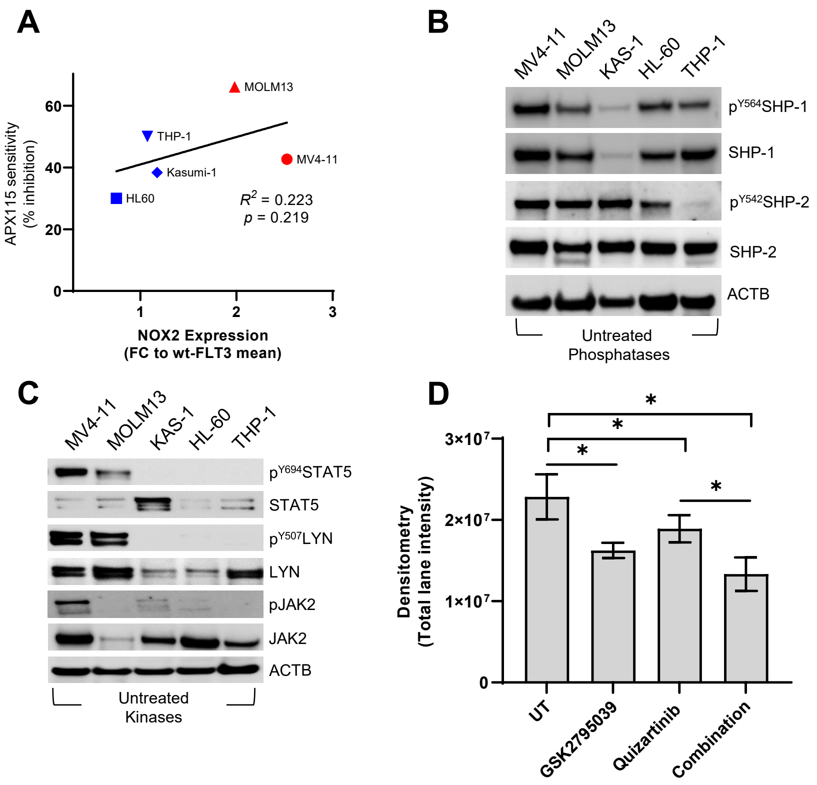


**Supplementary Figure S3.**

A

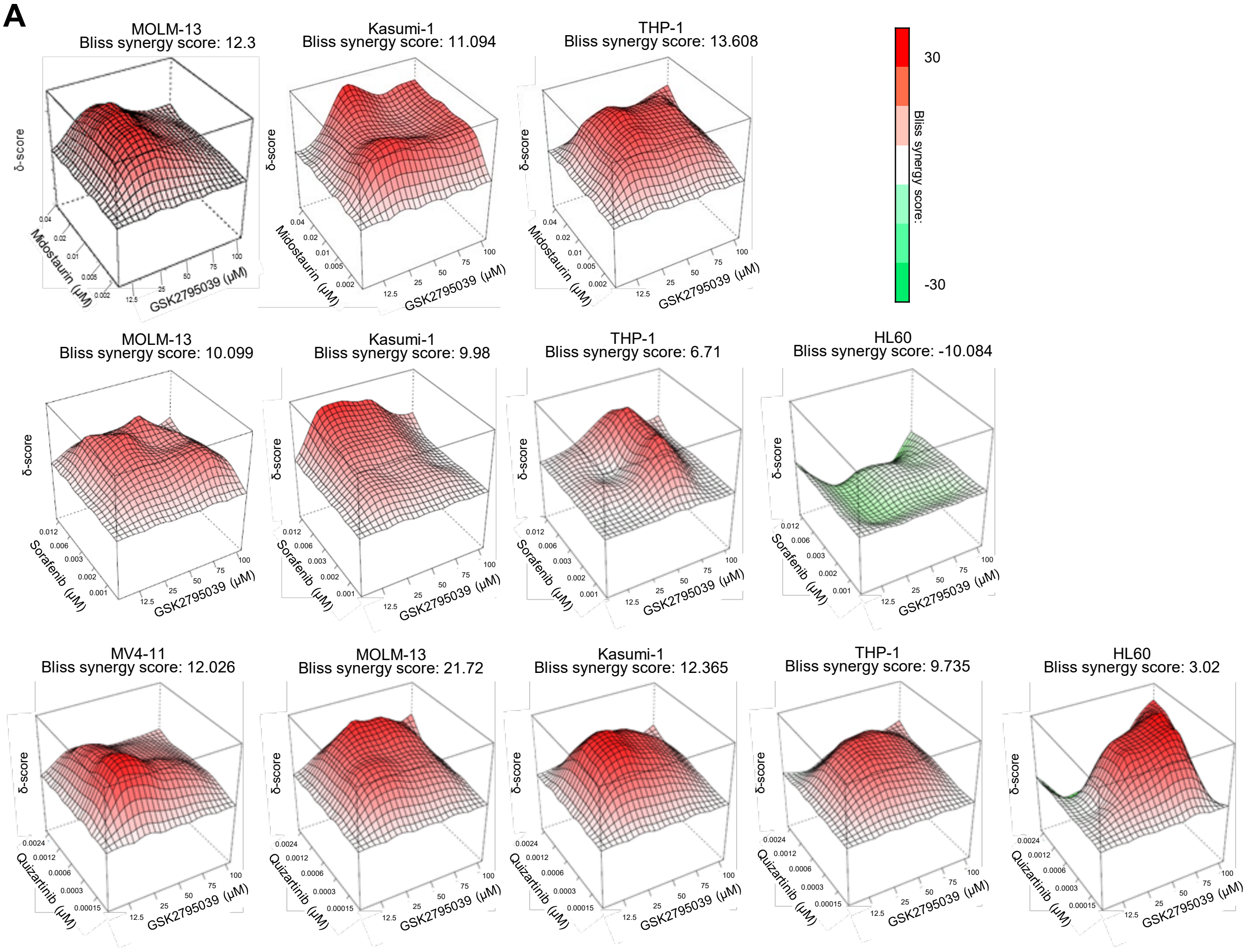


B

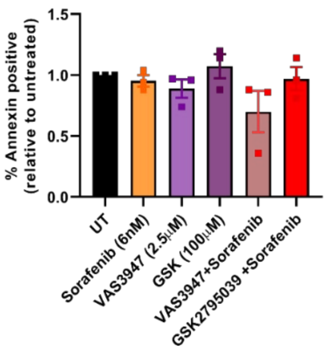


**Supplementary Figure S4.**

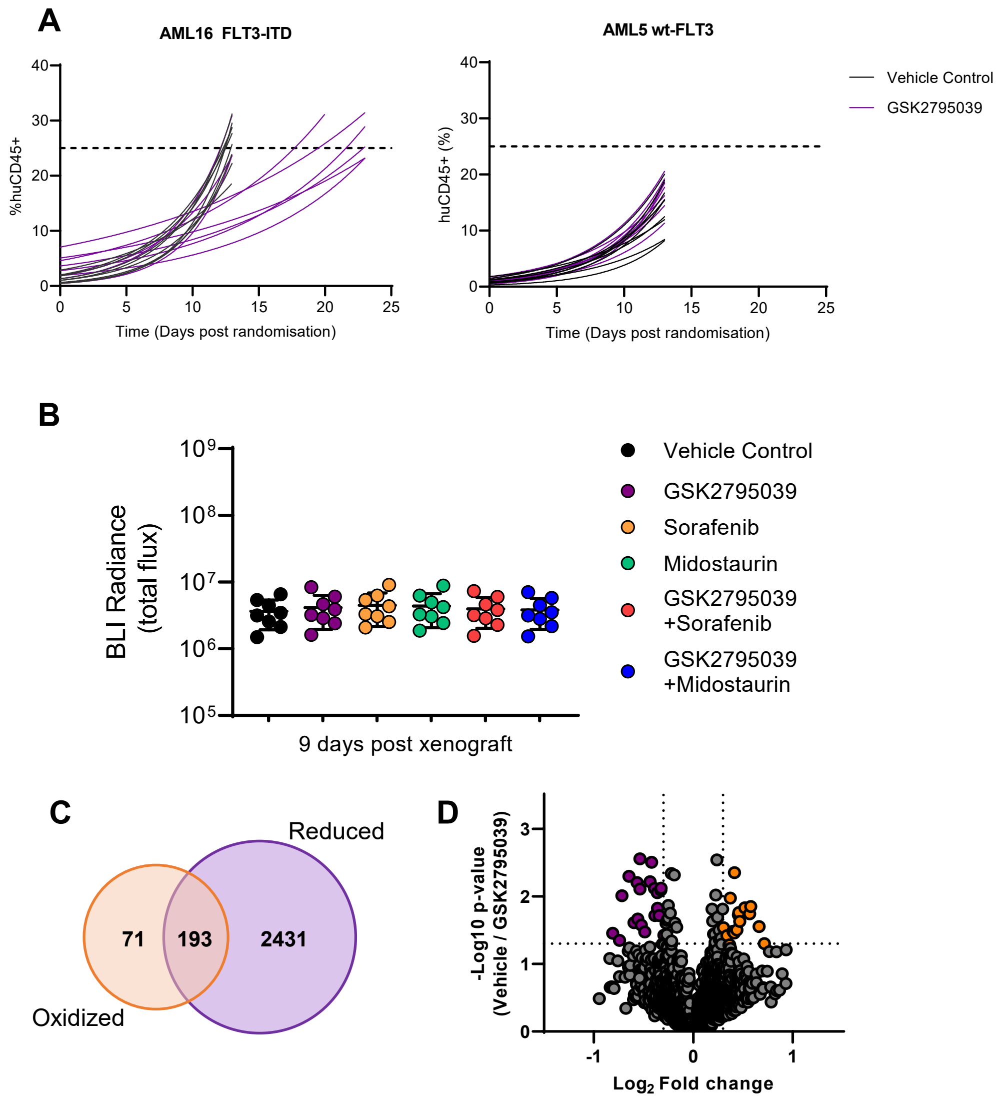


**Supplementary Figure S5.**
